# Translation efficiency covariation across cell types is a conserved organizing principle of mammalian transcriptomes

**DOI:** 10.1101/2024.08.11.607360

**Authors:** Yue Liu, Shilpa Rao, Ian Hoskins, Michael Geng, Qiuxia Zhao, Jonathan Chacko, Vighnesh Ghatpande, Kangsheng Qi, Logan Persyn, Jun Wang, Dinghai Zheng, Yochen Zhong, Dayea Park, Elif Sarinay Cenik, Vikram Agarwal, Hakan Ozadam, Can Cenik

## Abstract

Characterizing shared patterns of RNA expression between genes across conditions has led to the discovery of regulatory networks and novel biological functions. However, it is unclear if such coordination extends to translation. Here, we uniformly analyzed 3,819 ribosome profiling datasets from 117 human and 94 mouse tissues and cell lines. We introduce the concept of *<u>translation</u> <u>efficiency covariation</u>* (TEC), identifying coordinated translation patterns across cell types. We nominate candidate mechanisms driving shared patterns of translation regulation. TEC is conserved across human and mouse cells and uncovers gene functions that are not evident from RNA or protein co-expression. Moreover, our observations indicate that proteins that physically interact are highly enriched for positive covariation at both translational and transcriptional levels. Our findings establish TEC as a conserved organizing principle of mammalian transcriptomes.

## INTRODUCTION

Over the past three decades, technological advances have progressively revealed the expression of RNAs with increasing spatial and cellular resolution^1–7^. These measurements have driven conceptual advances including the widespread use of RNA co-expression analysis, which quantifies the similarity of gene expression patterns across biological conditions^8–11^. Such analyses have proved informative for inferring gene functions^9,12,13^, predicting protein-protein interactions^14,15^, and identifying shared regulatory mechanisms via common transcription factor binding sites^16,17^.

These findings suggest that RNA co-expression may serve as a proxy for the proteomic organization of cells. Quantification of protein abundance across numerous cell types and conditions has recently allowed this assumption to be explicitly tested. Surprisingly, coordinated patterns of protein abundance patterns are frequently not detected at the RNA level^18,19^, and physically interacting proteins are much more likely to have coordinated protein abundance than RNA co-expression^18–20^. These discrepancies suggest that post-transcriptional regulation likely plays a significant role in proteome organization.

Translation regulation may bridge this gap, yet whether translational efficiency is coordinated among functionally related genes across biological contexts remains an open question. There are three lines of evidence that suggest its plausibility. First, mammalian mRNAs bind various proteins to form ribonucleoproteins that influence their lifecycle from export to translation^21^. The set of proteins interacting with an mRNA varies with time and context, significantly altering the duration, efficiency, and localization of protein production. These observations led to the proposal of the post-transcriptional RNA regulon model, positing that functionally related mRNAs are regulated together post-transcriptionally^22,23^. Second, in *E. coli* and yeast, proteins within multiprotein complexes are synthesized in stoichiometric ratios^24,25^. However, in humans, evidence of such proportional synthesis is reported for only two complexes: ribosomes^25,26^ and the oxidative phosphorylation machinery^27^ in a limited number of cell lines. Third, the formation of many protein complexes is facilitated by the co-translational folding of nascent peptides^24,25,28,29,30^.

Despite these precedents, a general framework for detecting translation-level coordination across cell types has been lacking. Here, we analyzed thousands of matched ribosome profiling and RNA-seq datasets from more than 140 human and mouse cell lines and tissues. To quantify the similarity in TE patterns across cell types, we introduce the concept of *Translation Efficiency Covariation* (TEC). Analogous to RNA co-expression, TEC is common among functionally related genes. We demonstrate that TEC nominates novel gene functions, is enriched among physically interacting proteins, and is conserved across species. Together, these findings suggest TEC is a functionally relevant and evolutionarily conserved organizing principle of mammalian gene expression.

## RESULTS

### Integrated analysis of thousands of ribosome profiling and RNA-seq measurements enable quantitative assessment of data quality

We undertook a comprehensive, large-scale meta-analysis of ribosome profiling data to quantify TE across different cell lines and tissues. We collected 2,195 ribosome profiling datasets for humans and 1,624 experiments for mice, along with their metadata (Fig. 1a; Methods). Given that metadata is frequently reported in an unstructured manner and lacks a formal verification step, we conducted a manual curation process to rectify inaccuracies and collect missing information, such as experimental conditions and cell types used in experiments. One crucial aspect of our manual curation was pairing between ribosome profiling and corresponding RNA-seq when possible. Overall, 1,282 (58.4%) human and 995 (61.3%) mouse ribosome profiling samples were matched with corresponding RNA-seq data (table S1). The resulting curated metadata facilitated the uniform processing of ribosome profiling and corresponding RNA-seq data using an open-source pipeline^31^. We call the resulting repository harboring these processed files RiboBase (table S1).

**Fig. 1.**
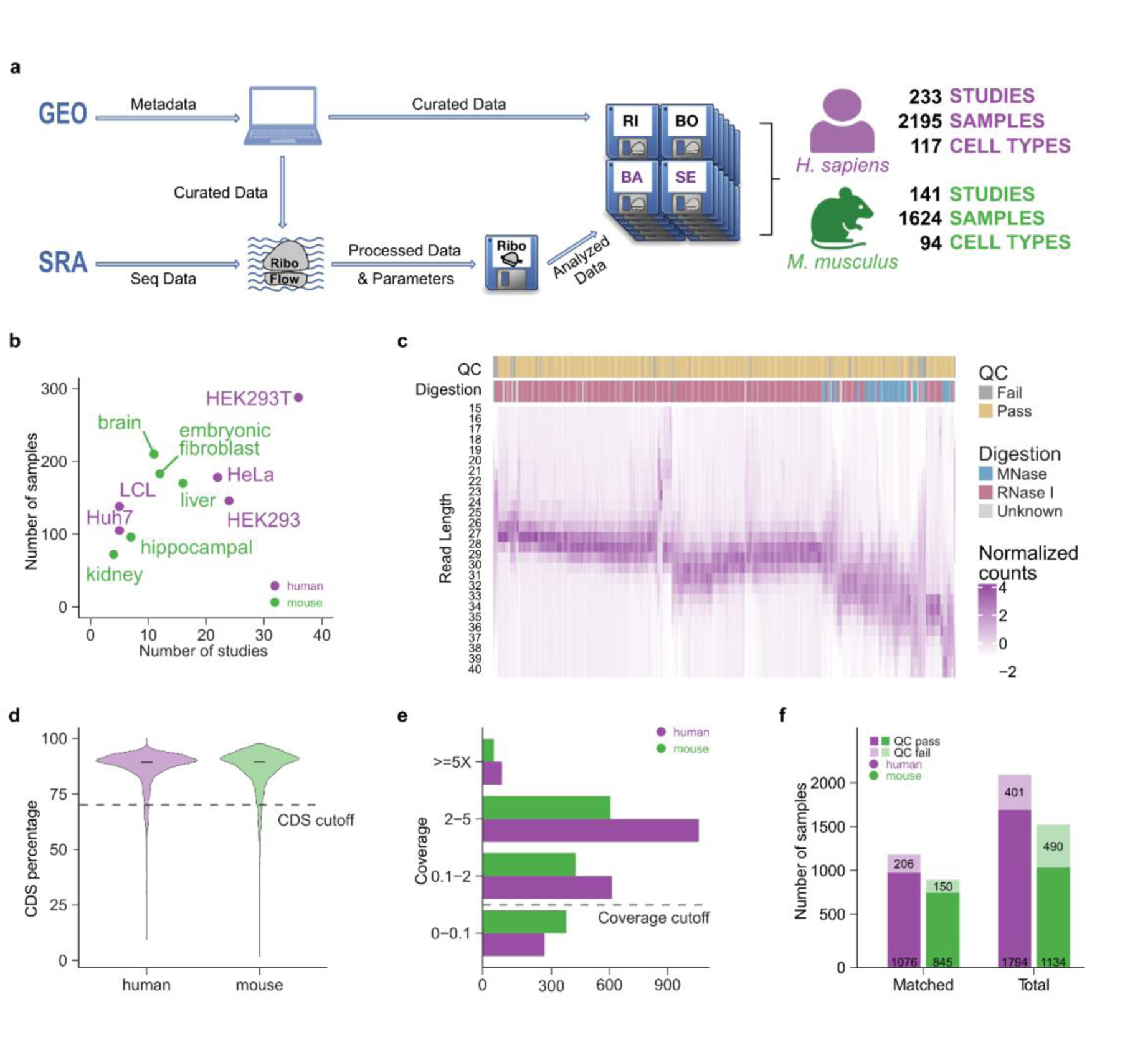
RiboBase: a comprehensive ribosome profiling database with thousands of experiments. **a,** Schematic of RiboBase. We manually curated metadata and processed the sequencing reads using a uniform pipeline (RiboFlow^31^). **b,** Top five most highly represented cell lines or tissues with respect to the number of experiments were plotted. **c,** We determined the ribonuclease used to generate ribosome profiling data for 680 experiments using human cancer cell lines. For each experiment, the read length distribution of RPFs mapping to coding regions was visualized as a heatmap. The color represents the z-score adjusted RPF counts (Methods). Each experiment where the percentage of RPFs mapping to CDS was greater than 70% and achieving sufficient coverage of the transcript (>= 0.1X) was annotated as QC-pass (Methods). **d,** For the 3,819 ribosome profiling experiments in RiboBase, we applied a function to select the range of RPFs for further analysis (Methods). We calculated the proportion of the selected RPFs that map to the coding regions (y-axis). The horizontal line represents the median of the distribution. **e,** Experiments (x-axis) were grouped by the transcript coverage (y-axis). **f,** Among the ribosome profiling experiments in RiboBase, 2,277 of them had corresponding RNA-seq data (matched). The number of samples that pass quality controls was plotted.

In RiboBase, the top cell types with the most experiments were HEK293T (13.1%) and HeLa (8.1%) for human; in mouse, the leading tissues were brain (9.6%), embryonic fibroblasts (8.3%), and liver (7.7%) (Fig. 1b; table S1). The median number of sequencing reads for ribosome profiling samples was ∼43.2 million for humans and ∼37.5 million for mice, respectively (Supplementary Fig. 1a-b; table S2-3; supplementary text). A majority of reads contained adapter sequences included during library preparation (with medians of 82.2% and 79.2% of total reads having adapters for human and mouse, respectively). Due to the substantial presence of ribosomal RNA in ribosome profiling datasets, only around 15% of total reads aligned to the transcript reference (Supplementary Fig. 1c-d; table S4-5; supplementary text).

The length of ribosome-protected mRNA footprints (RPFs) provides valuable information about data quality, the experimental protocol used, and translational activity^32^. The choice of nuclease impacts the resulting read length distribution of RPFs^33^ (Supplementary Fig. 2a-b). In agreement, we found that the peak position and range of RPF lengths were closely associated with the type of digestion enzymes used in human cancer samples (Fig. 1c). To account for the variability of RPF length distributions across the compendium of experiments, we developed a module that allowed for setting sample-specific RPF read length cutoffs (Supplementary Fig. 3a; Methods). This dynamic approach proved more effective than using fixed minimum and maximum values for RPF lengths, resulting in a higher retrieval of usable reads (median increase of 10.8% for human and 17.1% for mouse) and an increased proportion of reads within the coding sequence (CDS) region (Supplementary Fig. 3b).

After selecting a set of RPFs, we assessed the quality of ribosome profiling data within RiboBase using two additional criteria. Given that translating ribosomes should be highly enriched in annotated coding regions, we require that at least 70% of RPFs should be mapped to the CDS. We found that 160 human and 115 mouse samples failed to meet this criterion (Fig. 1d; table S6-7). Subsequently, we required a minimum number of RPFs that map to CDS to ensure sufficient coverage of translated genes (Methods). There were 318 human and 431 mouse samples with less than 0.1X transcript coverage (Fig. 1e; table S6-7). Altogether, 1,794 human samples and 1,134 mouse samples were retained for in-depth analysis. Of these, 1,076 human and 845 mouse samples were paired with matching RNA-seq data. Our results indicate a considerable fraction of publicly available ribosome profiling experiments had suboptimal quality (18.3% of the human and 30.1% of the mouse samples) (Fig. 1f). Interestingly, the data quality appeared to be independent of time (Supplementary Fig. 4). Additionally, we found that samples that passed our quality thresholds were more likely to exhibit three-nucleotide periodicity compared to those that failed quality control (92.59% vs 78.30% for humans and 91.36% vs 86.73% for mice; Supplementary Fig. 5; Methods). These findings underscore the necessity of meticulous quality control for the selection of experiments to enable large-scale data analyses.

### Translation efficiency is conserved across species and is cell-type specific

Ribosome profiling measures ribosome occupancy, a variable influenced by both RNA expression and translation. Thus, estimating translation efficiency necessitates analysis of paired RNA-seq and ribosome profiling data. To assess accurate matching in RiboBase, we first compared the coefficient of determination (R^2^) between matched ribosome profiling and RNA-seq data to that from other pairings within the same study. As would be expected from correct matching, we found that matched samples had significantly higher similarity on average (Fig. 2a; Welch two-sided t- test p-value < 2.2 x 10^-16^ for human and p-value = 2.1 x 10^-5^ for mouse). We then implemented a scoring system to quantitatively evaluate the correctness of our manual matching information (Methods). 99.2% of human samples and 98.5% of mouse samples had a sufficiently high matching score, demonstrating the effectiveness of our manual curation strategy (Supplementary Fig. 6a; Methods).

**Fig. 2.**
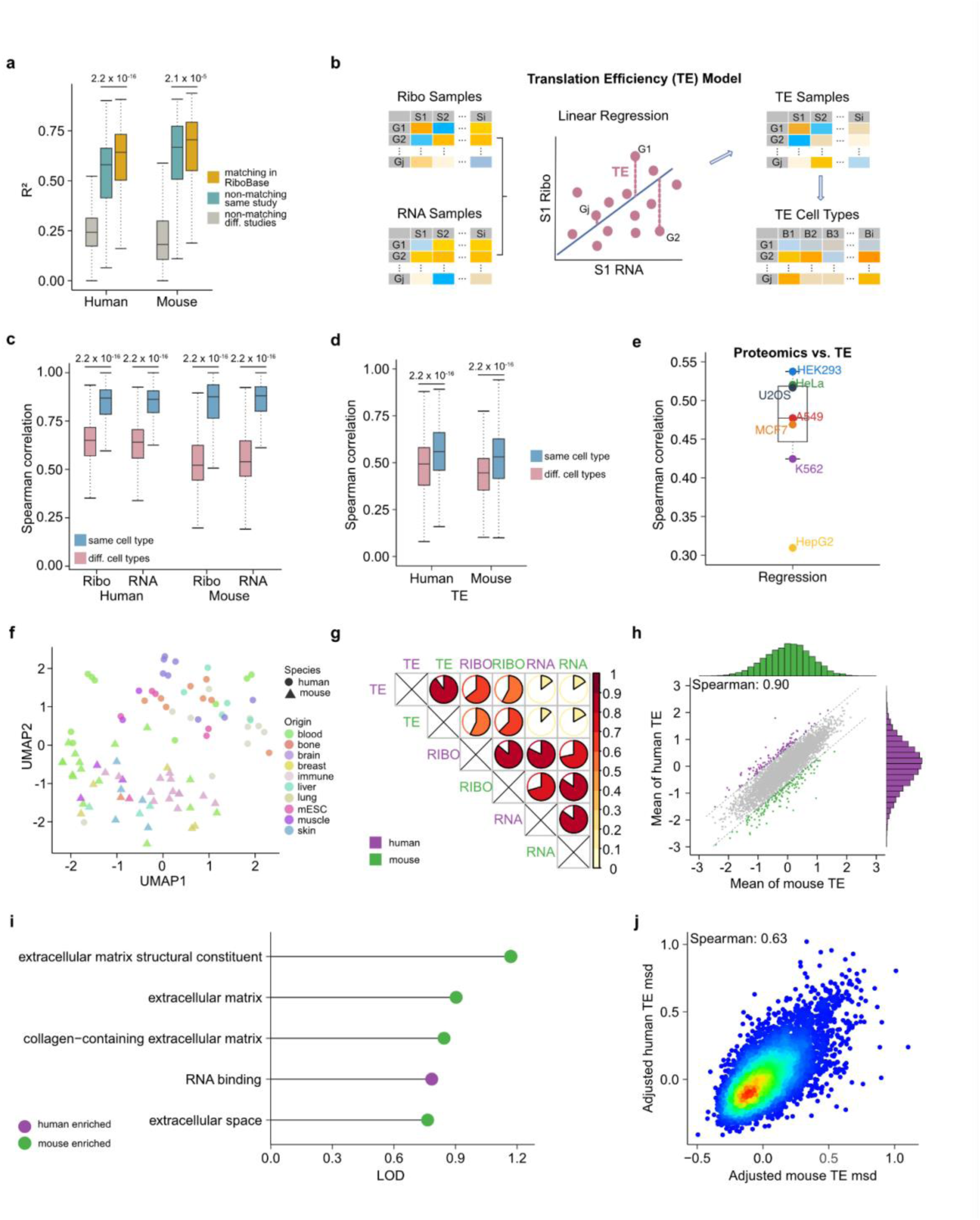
TE defined using a compositional linear regression model is conserved across cell types and species. **a,** The distribution of coefficient of determination (R², y-axis) between ribosome profiling data and RNA-seq in RiboBase was compared to random matching within the same study and across different studies. In each figure panel containing boxplots, the horizontal line corresponds to the median. The box represents the interquartile range (IQR) and the whiskers extend to the largest value within 1.5 times the IQR. The significant p-value shown in this figure was calculated using the two-sided Wilcoxon test. **b,** Schematic of TE calculation using the linear regression model with compositional data (CLR transformed; Methods; Supplementary Fig. 7). **c,** Distribution of correlations of TE (linear regression model) across experiments. **d,** The distribution of Spearman correlations between experiments (y-axis) was calculated based on whether they originated from identical or different cell lines or tissues. **e,** Correlation between TE and protein abundance from seven human cell lines^44^ was calculated using compositional regression method. **f,** We used UMAP to cluster the TE values of all genes across different cell types, considering only those origins with at least five distinct cell types. **g,** The Spearman correlation of 9,194 orthologous genes between human and mouse across TE, ribosome profiling, and RNA-seq levels. The circles represent the value of the Spearman correlation between groups. **h,** TE values were averaged across cell types and tissues for either human and mouse. Each dot represents a gene, and a 95% prediction interval was plotted to identify outlier genes (highlighted in purple and green). **i,** We conducted GO term enrichment analysis for outlier genes from panel H. We ranked the GO terms (y-axis) by the logarithm of the odds (LOD; x-axis). **j,** The correlation of the standard deviation of TE (quantified with adjusted metric standard deviation (msd); Methods; Supplementary Fig. 12c- d) for orthologous genes across different cell types between human and mouse.

Using the set of matched ribosome profiling and RNA-seq experiments, we next quantified TE, which is typically defined as the ratio of ribosome footprints to RNA-seq reads, normalized as counts per million^34^. However, this approach leads to biased estimates with significant drawbacks^35^. To address this limitation, we calculate TE based on a regression model using a compositional data analysis method^36–38^, avoiding the mathematical shortcomings of using the canonical log-ratio method (Fig. 2b; Supplementary Fig. 6a-c, 7; table S8-11; Methods; supplementary text).

We next assessed whether measurement errors due to differences in experimental procedures dominate variability that would otherwise be attributed to biological variables of interest. Specifically, we compared similarities between experiments that used the same cell type or tissue in different studies (Supplementary Fig. 8a). We found that ribosome profiling or RNA experiments from the same cell type or tissue exhibited higher similarity compared to those from different cell lines or tissues (Fig. 2c). Consistent with this observation, TE values displayed higher Spearman correlation coefficient within the same cell type or tissue (median correlation coefficient of 0.56 and 0.53 in human and mouse, respectively) compared to different cell lines and tissues (median correlation coefficient of 0.49 and 0.45 in human and mouse, respectively) (Fig. 2d).

We expected that an accurate estimate of TE would correlate significantly with protein abundance. We calculated for each transcript the cell type-specific TE by taking the average of TE values across all experiments conducted with that particular cell line and found that TE derived using the regression approach with winsorized read counts (Supplementary Fig. 8b; Supplementary Fig. 9- 11; supplementary text) was significantly correlated with protein abundance in seven cancer cell lines (mean Spearman correlation coefficient of 0.465; Fig. 2e).

Furthermore, TE measurements from cell lines and tissues with the same biological origin (e.g., blood) tended to cluster together, supporting the existence of cell-type-specific in addition to species-specific differences in TE (Fig. 2f). As expected, mean ribosome occupancy and RNA expression across cell types showed a strong correlation (Spearman correlation: ∼0.8), yet mean TE was only weakly associated with RNA expression (Spearman correlation: ∼0.2) (Fig. 2g).

Measurements of TE in two species across a large number of cell types enabled us to investigate the conservation of TE, ribosome occupancy, and RNA expression. Transcriptomes, ribosome occupancy, and proteomes exhibit a high degree of conservation across diverse organisms^39,40^. Consistently, we found average ribosome occupancy, RNA expression, and TE across different cell lines and tissues were highly similar between orthologous genes in human and mouse (Fig. 2g; table S12). Specifically, the Spearman correlation coefficient of mean TE across cell types and tissues between human and mouse was 0.9 (Fig. 2h), which is comparable to the mean RNA expression correlation between human and mouse (∼0.86, Supplementary Fig. 12a). Using a 95% prediction interval to identify outlier genes, we found that outlier genes with higher mean TE in humans compared to mice were enriched in the gene ontology term ‘RNA binding function’ (Fig. 2i). In contrast, genes with elevated mean TE in mice were enriched for having functions related to extracellular matrix and collagen-containing components (Fig. 2i). The enrichment of genes with higher TE in mice, particularly those from the extracellular matrix and collagen-containing components, may be due to the fact that many samples in mouse studies are derived from the early developmental stage^41^.

Despite the high correlation of mean TE across various cell lines and tissues between human and mouse, TE distinctly exhibits cell-type specificity. While several studies compared the conservation of TE between the same tissues of mammalians or model organisms^40,42,43^, our dataset uniquely enabled us to determine the conservation of variability of TE for transcripts across different cell types. Intriguingly, we observed a moderately high similarity between the variability of TE of orthologous genes in human and mouse (Spearman partial correlation coefficient = 0.63; Fig. 2j; Supplementary Fig. 12b-d; Methods). Our results reveal that certain genes exhibit higher variability of TE across cell types and this is a conserved property between human and mouse.

### Translation efficiency covariation (TEC) is conserved between human and mouse

Uniform quantification of TE enabled us to investigate the similarities in TE patterns across cell types. Given the usefulness of RNA co-expression in identifying shared regulation and biological functions, we aimed to establish an analogous method to detect patterns of translation efficiency similarity among genes. To achieve this, we employed the proportionality score (rho)^37,38^, a statistical method that quantifies the consistency of how relative TE changes across different contexts (Methods). Recent work suggested that the proportionality score enhances cluster identification in high-dimensional single-cell RNA co-expression data^10^. Consistent with these findings, our analysis revealed its particular effectiveness in quantifying ribosome occupancy covariation (Supplementary Fig. 13; Methods). We calculated rho scores for all pairs of human or mouse genes where a high absolute rho score indicates significant translation efficiency covariation (TEC) between pairs (Fig. 3a).

**Fig. 3.**
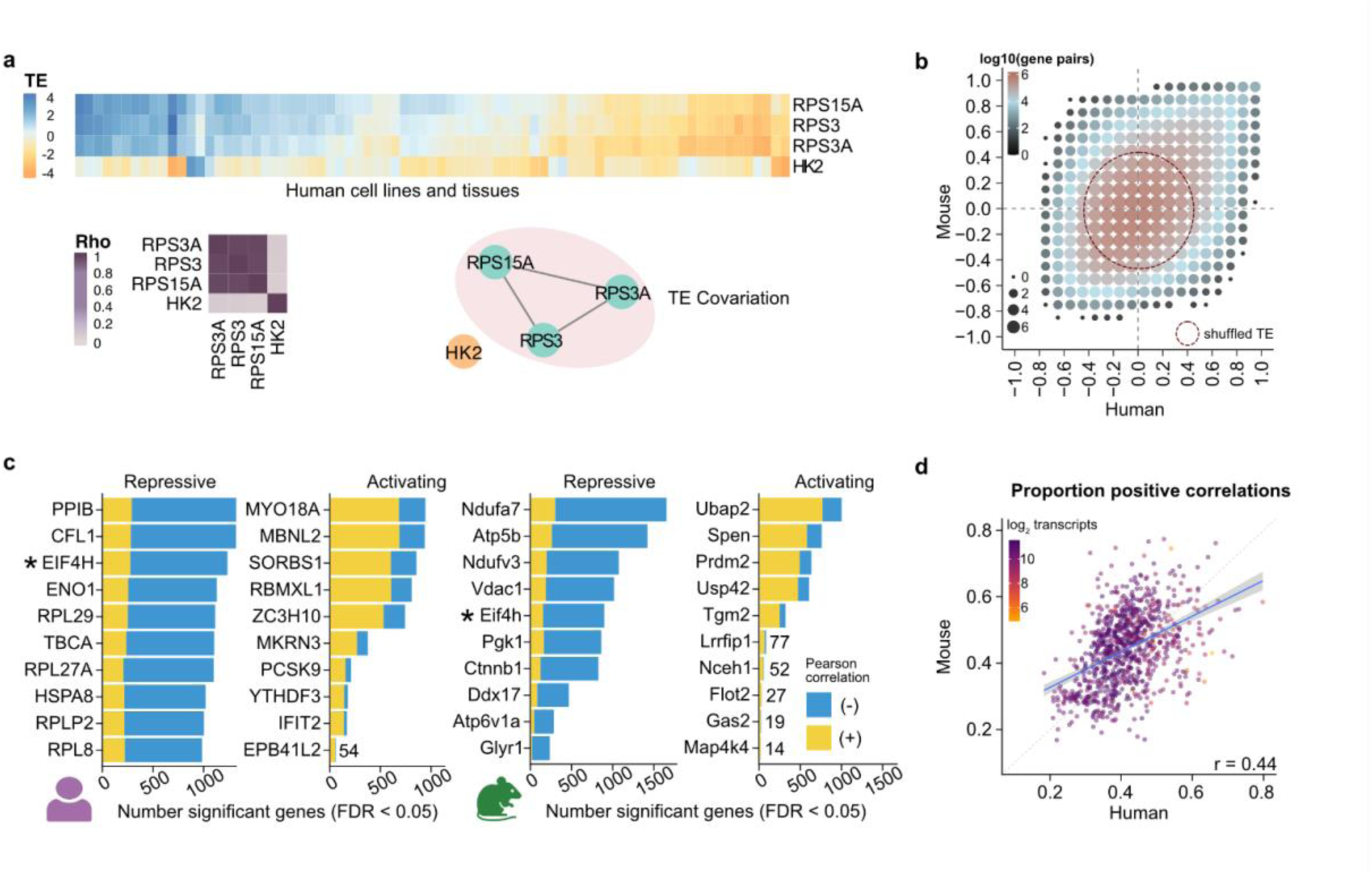
Translation efficiency covariation is conserved between human and mouse. **a,** Example illustrating TEC between genes. The top section presents TE patterns across cell types in human. The bottom left part displays the similarity of the pattern between these genes quantified using proportionality scores. **b,** We calculated the TEC for gene pairs and compared their differences for the same orthologous gene pairs between human and mouse. In the figure panel, each dot size represents the aggregated log10-transformed counts of gene pairs falling within specified ranges. We also calculated TEC using randomized TE for each gene (shuffled). The red dashed line in the figure captures the 95% gene pair TEC values obtained with shuffled TE (Supplementary Fig. 14). **c,** The top ten RBPs with the highest number of genes showing significant correlations between transcript TE and RBP RNA expression are displayed. An asterisk marks genes in the top ten in both species. **d,** For all RBPs (each dot), we plotted the proportion of positive correlations between TE of significant transcripts and the RNA expression of the RBP. Blue line is a linear fit with 95% confidence intervals in gray. Pearson correlation coefficient is shown.

Previous studies have indicated that RNA co-expression between genes is conserved in mammals^39,45,46^. To assess the potential evolutionary significance of the newly introduced TEC concept, we evaluated its conservation across human and mouse transcripts. Indeed, TEC was highly similar for orthologous gene pairs in humans and mice (Fig. 3b, Pearson correlation coefficient 0.41), compared to a negligible correlation in TEC derived from shuffled TE values (Supplementary Fig. 14, Pearson correlation coefficient 0.00022). Our findings imply that translation efficiency patterns are evolutionarily preserved, paralleling the conservation of RNA co-expression.

There are few known examples of transcript sets with shared translational control across cell types, including those with 5′ terminal oligopyrimidine (TOP) motifs^47^ and transcripts regulated by CSDE1^48^. We hypothesized that these sets of transcripts would exhibit significantly higher TEC compared to a control set, and this was indeed observed (Supplementary Fig. 15a; Wilcoxon p-value < 2.2e-16). These findings highlight that TEC effectively captures previously established patterns of shared translational regulation among transcripts.

RNA co-expression analyses led to the discovery of regulatory motifs and shared transcription factor binding sites^49^. We hypothesized that TEC among genes may similarly nominate RNA binding proteins (RBPs) as potential drivers of TEC^50^. We identified groups of transcripts whose TE is correlated with the RNA expression of RBPs^51^ (1274 human and 1762 mouse RBPs; Methods; https://zenodo.org/uploads/11359114). For example, the RNA expression of ZC3H10 has largely positive correlations with TE of transcripts (71% of all significant correlations Fig. 3c). Conversely, the RNA expression of subunits of ubiquinone oxidoreductase (Ndufa7, Ndufv3) is negatively correlated with TE for many transcripts in mice (Fig. 3c).

We observed a correspondence between human and mouse in the proportion of transcripts whose TE is significantly correlated with an RBP’s expression (Pearson correlation 0.44; Fig. 3d). However, knockout of three candidate RBPs did not lead to significant changes in TE (Supplementary Fig. 16), which may be due to several limitations including cell-type specificity, efficiency of knockout, and limited replicates (supplementary text). Taken together, our analyses nominate RBPs that may coordinate the TEC of evolutionarily conserved networks.

### TEC between transcripts across cell lines and tissues is associated with shared biological functions

Given that co-expression at the RNA level is predictive of shared biological functions^11,52,53^, we next assessed whether TEC indicates common biological roles among genes. We calculated the area under the receiver operating characteristic curve (AUROC) to measure the ability of TEC in distinguishing genes with the same biological functions (Methods). Genes that are annotated with a common GO term exhibited a similar degree of RNA co-expression and TEC, both of which were significantly higher than would be expected by chance (Median AUROC across GO terms calculated with TEC: 0.63 for human, 0.65 for mouse; RNA co-expression RNA: 0.66 for human, 0.69 for mouse; Fig. 4a; table S13-14; Methods). These findings demonstrate that TEC, similar to RNA co-expression, serves as an indicator of shared biological functions among genes.

**Fig. 4.**
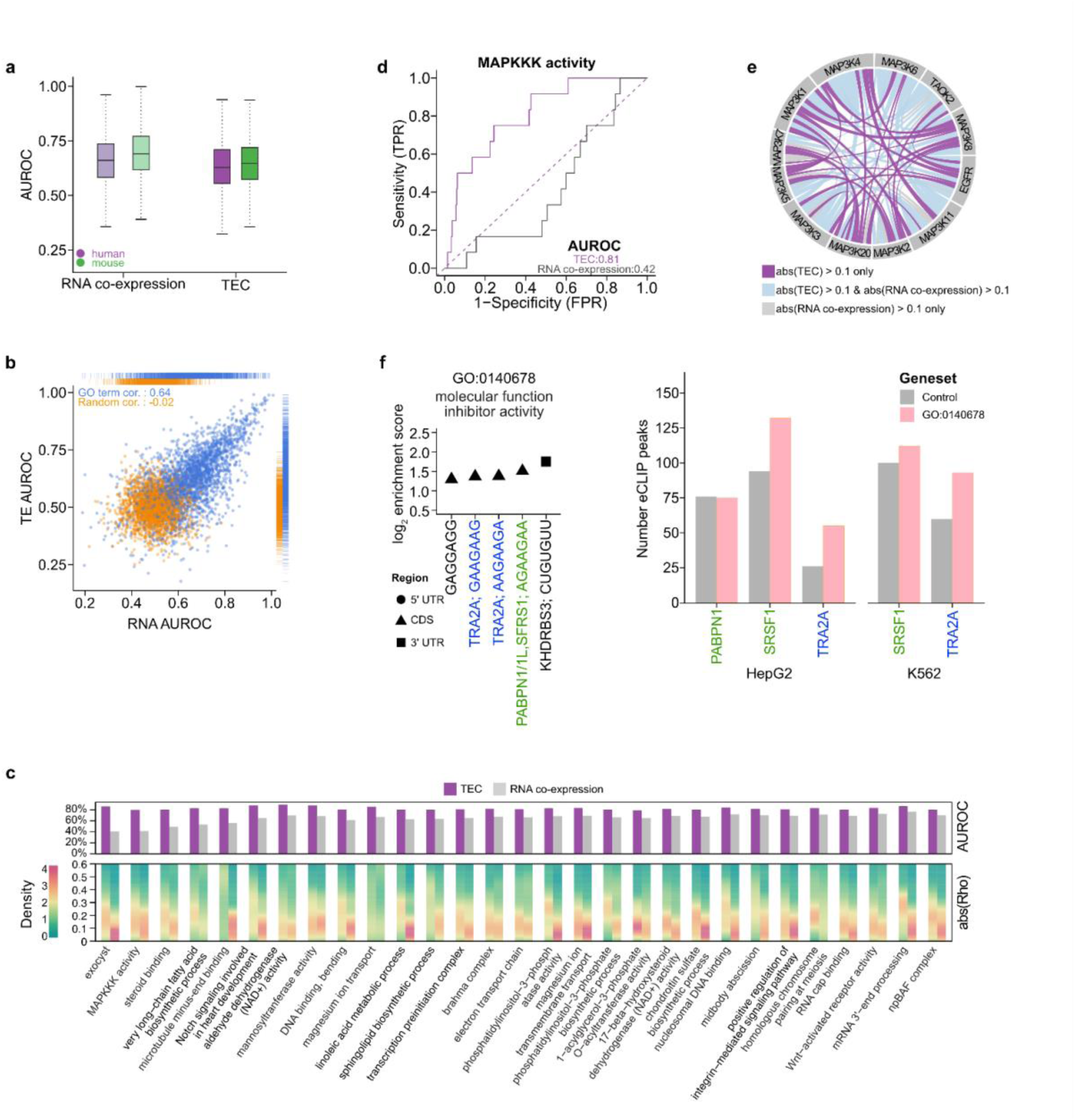
Genes associated with certain biological functions exhibit higher similarity patterns in TE than in RNA expression. **a,** We calculated the similarity of expression (quantified by AUROC; y-axis) among genes within 2,989 human and 3,340 mouse GO terms. In the box plot, the horizontal line corresponds to the median. The box represents the IQR and the whiskers extend to the largest value within 1.5 times the IQR. **b,** Each blue dot represents the AUROC calculated for a given GO term using TEC and RNA co-expression levels. Orange dots represent the same values for random grouping of genes (Methods). **c,** For GO terms where genes exhibit greater similarity at the TE level than at the RNA expression level (AUROC for TEC > 0.8, and difference of AUROC measured with TEC and RNA co-expression > 0.1), we visualized the distribution of absolute rho scores for gene pairs (bottom; gene pairs with abs(rho) > 0.1). **d,** AUROC plot calculated with genes associated with MAPKKK activity. **e,** In the circle plot, the connections display absolute rho above 0.1 either at TE level alone (purple), at both RNA and TE levels (blue), or RNA level alone (gray) for gene pairs involved in MAPKKK activity. **f,** Motif enrichment (left) for the GO term ‘molecular function inhibitor activity’ (Supplementary Fig. 17b). RNA binding proteins (RBPs) matching the motifs from oRNAment^59^ or Transite^60^ are indicated. Enhanced cross-linking immunoprecipitation (eCLIP) data^61^ indicates increased binding of TRA2A and SRSF1 in the CDS of genes for this GO term compared to matched control genes with similar sequence properties (Methods).

Furthermore, we observed that biological functions whose members exhibit a high degree of RNA co-expression were also likely to have TEC. Specifically, the Spearman correlation between the AUROC scores calculated using TEC and RNA co-expression was ∼0.64 for human GO terms in contrast to ∼-0.02 when random genes were grouped (Fig. 4b). Despite the low correlation between average RNA expression and TE for human genes (Fig. 2g), our results highlight that members of specific biological functions whose RNA expression is coordinated across cell types tend to exhibit consistent translation efficiency patterns. This finding suggests coordinated regulation at both transcriptional and translational levels among functionally related genes.

While many gene functions were predicted accurately with both RNA co-expression and TEC, we noted specific exceptions. Notably, genes in 29 human GO terms demonstrated significantly stronger TEC than RNA co-expression (at least 0.1 higher AUROC; Fig. 4c; Supplementary Fig. 17-18; supplementary text). An example of such a GO term is ‘MAPKKK activity’ (Fig. 4d-e). While there is limited evidence of direct translational regulation of the MAPKKK family, the RBP IMP3 may provide a potential mechanism for such regulation^54^. These results indicate that some genes with specific biological functions exhibit greater similarity at the translational level.

We hypothesized that genes with shared functions and high TEC may be regulated through a common mechanism, analogous to shared transcription factor binding sites that mediate RNA co- expression^55,56^. Accordingly, we expected these genes to harbor shared sequence elements. We identified enriched heptamers in the transcripts of five human and three mouse GO terms with significant TEC and at least 12 genes in the GO term (AUROC measured with TEC > 0.7, difference in AUROC between TEC and RNA co-expression > 0.2; Fig. 4f; Supplementary Fig. 17b; Supplementary Fig. 18e; Methods). For example, we found AG-rich motifs in coding regions of human genes with “molecular function inhibitor activity” (Fig. 4f). These motifs match the known binding sites of three RBPs (TRA2A, PABPN1, and SRSF1). In line with the enrichment of these motifs, analysis of eCLIP data revealed increased deposition of these RBPs in the coding sequences of genes in this GO term compared to matched control transcripts (Fig. 4f; Methods). Furthermore, we identified several additional enriched heptamers that currently have no RBP annotations, suggesting these motifs might be targets for RBPs that have not yet been characterized (also see supplementary text for sequence features associated with TEC). In contrast, our analyses indicate miRNA is not a primary driver of TEC (Supplementary Fig. 19), in agreement with previous literature^57,58^.

### TEC nominates gene functions

We next investigated whether gene functions may be predicted by utilizing TEC, given the success of RNA co-expression for this task^52,53^. The functional annotations of human genes are continuously being updated, providing an opportunity to test this hypothesis using recently added information to the knowledge base. Specifically, we used functional annotations from the GO database from January 1, 2021, to determine functional groups that demonstrate strong TEC among its members (AUROC > 0.8) and developed a framework to predict new functional associations with these groups (Methods). By comparing our predictions to annotations from December 4, 2022, we confirmed the predicted association of the *LOX* gene with the GO term ‘collagen-containing extracellular matrix’. LOX critically facilitates the formation, development, maturation, and remodeling of the extracellular matrix by catalyzing the cross-linking of collagen fibers, thereby enhancing the structural integrity and stability of tissues^62,63^. Our prediction successfully identified this new addition, as *LOX* exhibits positive similarity in TE with the vast majority of genes in this term (Fig. 5a).

**Fig. 5.**
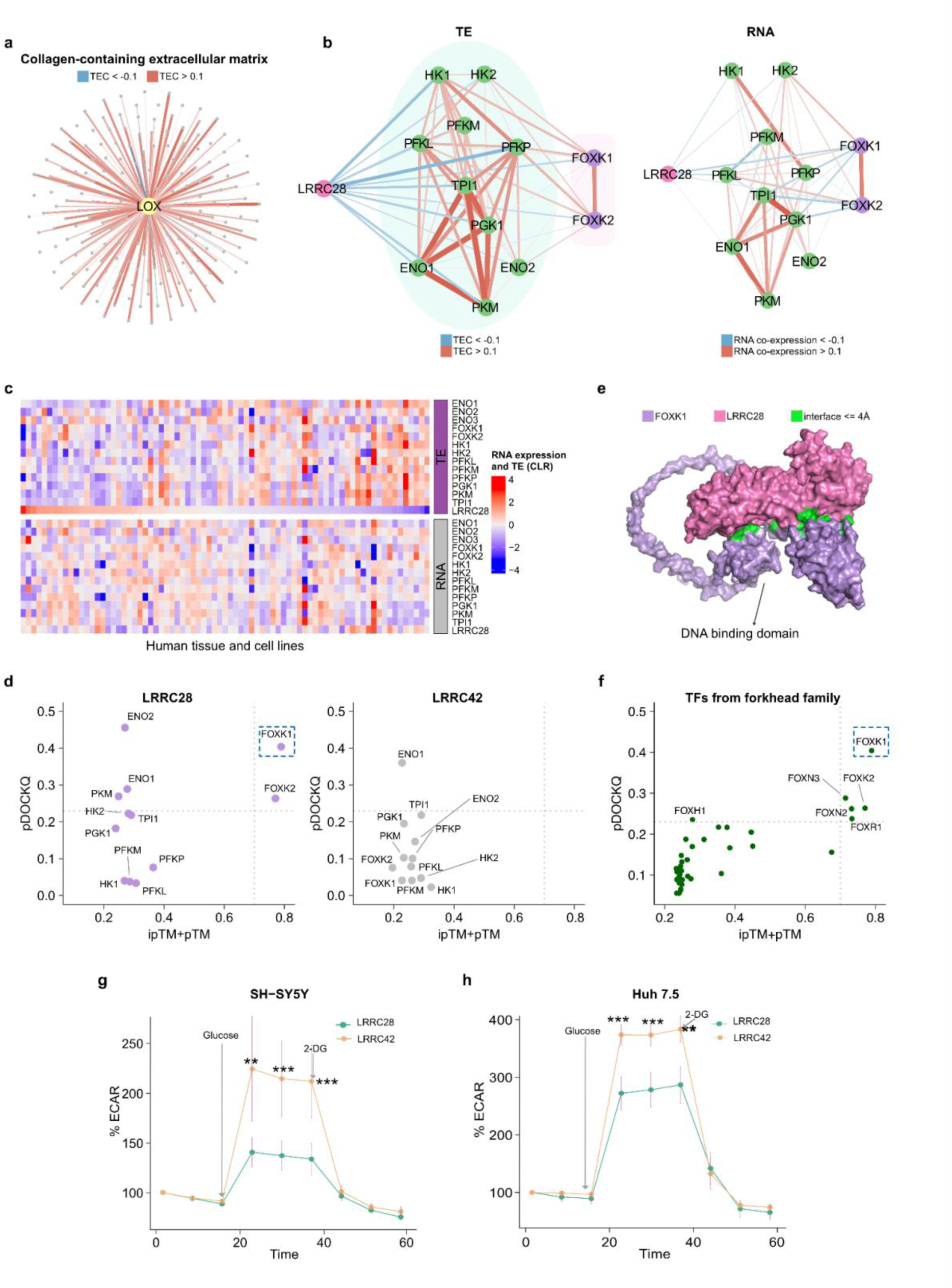
TEC enables the prediction of novel gene functions. **a,** We predicted that LOX belongs to the collagen-containing extracellular matrix using an older version of human GO terms (January 1, 2021) and confirmed this prediction with the newer version (December 4, 2022; Methods). The network displays the similarity in TE between LOX (yellow dot) and other genes (gray dots) from the collagen-containing extracellular matrix. Line weight in figure panels indicates the absolute value of rho from 0.1 to 1. **b,** The networks display the rho between LRRC28 and glycolytic genes at the TE level (on the left) and RNA level (on the right) in humans. Green dots represent genes that belong to the glycolysis pathway, purple nodes are transcription factors that regulate glycolysis. **c,** TE and RNA expression of *LRRC28*, glycolytic genes, and transcription factors regulating glycolysis (*FOXK1*, *FOXK2*) across human cell types and tissues. **d,** We used AlphaFold2-Multimer to calculate the binding probabilities between the proteins LRRC28 or LRRC42 and glycolytic proteins (Methods). We evaluated the models with ipTM+pTM (x-axis) and precision of protein-protein interface binding predictions (pDOCKQ; y-axis). We set a threshold of ipTM+pTM > 0.7^75^ and pDOCKQ > 0.23^76,77^ as previously suggested to identify confident binding. **e,** 3D model of binding between LRRC28 and FOXK1. For visualization purposes, we removed residues 1-101 and 370-733 in FOXK1 (pLDDT scores below 50). **f,** Binding probabilities between LRRC28 and transcription factors belonging to the forkhead family^78^. The dashed lines represent ipTM+pTM > 0.7 or pDOCKQ > 0.23. Kinetic ECAR response of **g,** SHSY-5Y cell line (n=6, stable overexpression) and **h,** Huh 7.5 cell line (n=6; transient overexpression) overexpressing LRRC28 or LRCC42 to 10 mM glucose and 100 mM 2- DG. Unpaired two-sided Student’s *t*-test, \*\*\**P* < 0.005, \*\**P* < 0.05. Panel g & h shows mean ± s.d.; *n* shows biological independent experiments.

Recognizing the capacity of TEC to elucidate biological functions, we utilized a recent version of GO annotations (December 4, 2022) to systematically predict new associations for genes. To underscore the unique insights gained from TEC, we focused on the 33 human and 31 mouse GO terms that either exhibited significantly higher TEC than RNA co-expression (Table 1) or provided new functional predictions that were only supported by TEC (the ranking of the newly predicted gene with RNA co-expression fell beyond the top 50%, table S15-16; Methods). By focusing on these GO terms, we aimed to identify similarity patterns based on TE, revealing functional associations that would not be detected by RNA co-expression. We conducted a literature search to determine if prior research supported these predictions, and found that 11 have already been corroborated by previous publications, although they have not yet been reflected in the relevant GO term annotations (Table 1; supplementary text). For example, cryo-electron microscopy experiments demonstrated that human DNMT1 binds to hemimethylated DNA in conjunction with ubiquitinated histone H3^64^. This binding facilitates the enzymatic activity of DNMT1 in maintaining genomic DNA methylation. Our analysis revealed that DNMT1 was the highest ranking prediction exhibiting strong TEC with genes associated with nucleosomal DNA binding function. In mouse, we predicted Plekha7 to be a member of the regulation of developmental processes. This prediction was recently validated by the observation of neural progenitor cell delamination upon the disruption of Plekha7^65–69^.

**Table 1:** Literature support for gene functions predicted using TEC. In the table, we list the predictions that are supported by literature. To do the new gene function prediction, we selected GO terms with AUROC measured with TEC >=0.8, then focused on the subset with differences between AUROC measured with TEC and RNA co-expression >= 0.1.

| Term | Species | Description | New adding gene (top ranking in TE) | TEC AUROC | RNA co-expression AUROC | Ranking of new adding gene in RNA | Reference |
| --- | --- | --- | --- | --- | --- | --- | --- |
| GO:0005496 | Human | steroid binding | EDNRA | 0.81 | 0.50 | 3991 | 135 |
| GO:0022900 | Human | electron transport chain | TMEM70 | 0.82 | 0.67 | 1611 | 136,137 |
| GO:0042813 | Human | Wnt-activated receptor activity | HSPG2 | 0.85 | 0.74 | 341 | 138,139 |
| GO:0007129 | Human | homologous chromosome pairing at meiosis | KIF4A | 0.84 | 0.73 | 11 | 140,141 |
| GO:0031492 | Human | nucleosomal DNA binding | DNMT1 | 0.85 | 0.73 | 251 | 64,142,143 |
| GO:0050793 | Mouse | regulation of developmental | Plekha7 | 0.85 | 0.70 | 3929 | 65–69 |

| **Term** | **Species** | **Description** | **New adding gene (top ranking in TE)** | **TEC AUROC** | **RNA co-expression AUROC** | **Ranking of new adding gene in RNA** | **Reference** |
| --- | --- | --- | --- | --- | --- | --- | --- |
|  |  | process |  |  |  |  |  |
| GO:1990023 | Mouse | mitotic spindle midzone | Cenpf | 0.90 | 0.74 | 54 | 144,145 |
| GO:0016342 | Mouse | catenin complex | Fat1 | 0.85 | 0.73 | 781 | 146–149 |
| GO:0005539 | Mouse | glycosaminoglycan binding | Lox | 0.95 | 0.83 | 32 | 150,151 |
| GO:0140374 | Mouse | antiviral innate immune response | Arhgap31 | 0.84 | 0.73 | 1085 | 152–156 |
| GO:0022010 | Mouse | Central nervous system myelination | Jam2 | 0.88 | 0.76 | 584 | 157 |

The high rate of validation of our predictions in the literature suggested that other predictions based on TEC may reflect new and yet to be confirmed functions. In particular, we observed that the human leucine-rich repeat-containing 28 (*LRRC28*) gene displays strong TEC with glycolytic genes, but is not co-expressed at the RNA level (Fig. 5b-c, table S17). Specifically, *LRRC28* displayed negatively correlated TE with key glycolytic genes including *HK1*, *HK2*, *PFKL*, *PFKM*, *PFKP*, *TPI1*, *PGK1*, *ENO1*, *ENO2*, *PKM*, and two transcription factors *FOXK1* and *FOXK2* that regulate glycolytic genes^70^. Given that the leucine-rich repeat domains typically facilitate protein- protein interactions^71^, LRRC28 may interact directly with one or more of the glycolytic proteins. Using AlphaFold2-Multimer^72^, we calculated the binding confidence scores between LRRC28 and all glycolysis-associated proteins (Methods) and found that LRRC28 has a very high likelihood of binding to FOXK1 (Fig. 5d-e).

FOXK1 is a member of the forkhead family of transcription factors that share a structurally similar DNA-binding domain^73,74^. Interestingly, LRRC28 likely binds both the non-DNA-binding region and DNA-binding domain of FOXK1 (distance < 4 angstroms; Fig. 5e; Supplementary Fig. 20a). This observation led us to examine the specificity of the interaction between LRRC28 and FOXK1. We calculated the binding probabilities of LRRC28 with 35 other forkhead family transcription factors, finding that FOXK1 exhibits the strongest evidence of physical interaction with LRRC28 (Fig. 5f). This specificity is potentially due to a unique binding site between LRRC28 and FOXK1’s non-DNA-binding region (Fig. 5e). As an additional control, we selected LRRC42, a protein with leucine-rich repeats that does not exhibit TEC with glycolytic genes. As expected, LRRC42 showed a very low likelihood of interaction with any of the glycolytic genes, including FOXK1 (Fig. 5d). These findings suggest that LRRC28 may serve as a regulator of glycolysis by binding to FOXK1, thereby preventing FOXK1 from binding to the promoter regions of glycolytic genes and leading to the downregulation of glycolysis. To experimentally test this prediction, we stably overexpressed LRRC28 in four human cell lines and assessed the glycolytic capacity of the cells by quantifying the extracellular acidification rate (ECAR). Stable overexpression of LRRC28 in Huh 7.5 resulted in slower growth therefore we resorted to transient overexpression strategy for this particular cell line. Congruent to our expectation, ECAR was found to be significantly lower in SH-SY5Y and Huh 7.5 cells overexpressing LRRC28 compared to cells overexpressing a control gene; LRRC42 (Fig. 5g-h). However, HEK293T and MCF-7 showed non-significant changes between the two conditions (Supplementary Fig. 20b-c). Interestingly, the varied response in presence of LRRC28 likely reflects the metabolic state differences between cell lines and suggests differential dependency on LRRC28. Taken together, TEC reveals shared biological functions and predicts novel associations, providing insights not attainable with RNA co-expression analysis alone.

### Genes with positive TEC are more likely to physically interact

The predicted binding between LRRC28 and FOXK1 suggests the utility of TEC to reveal physical interactions between proteins. Proteins that physically interact tend to be co-expressed at the RNA level^14,79,80^, and many protein complexes are assembled co-translationally^81^, leading us to hypothesize that the TE of interacting proteins may be coordinated across cell types. Specifically, we expect that there should be positive covariation between the TE of interacting proteins to ensure their coordinated production^24,25^. To test this hypothesis, we categorized gene pairs by whether they display positive or negative similarity in RNA expression or TE across cell types. Compared to all possible pairs (124,322,500), or those with the same biological function (6,492,564), physically interacting pairs of proteins (1,030,794 from STRING database^80^) were substantially enriched for positive similarity of TE and RNA expression patterns (Fig. 6a; chi-square test p< 2.2 x 10^-16^ and 1.88-fold enrichment compared to all pairs; table S18). This result aligns with the notion that genes with the same function can be regulated in opposite directions, as indicated by negative rho values, in contrast to physically-interacting proteins^82^ which are enriched for positive rho values.

**Fig. 6.**
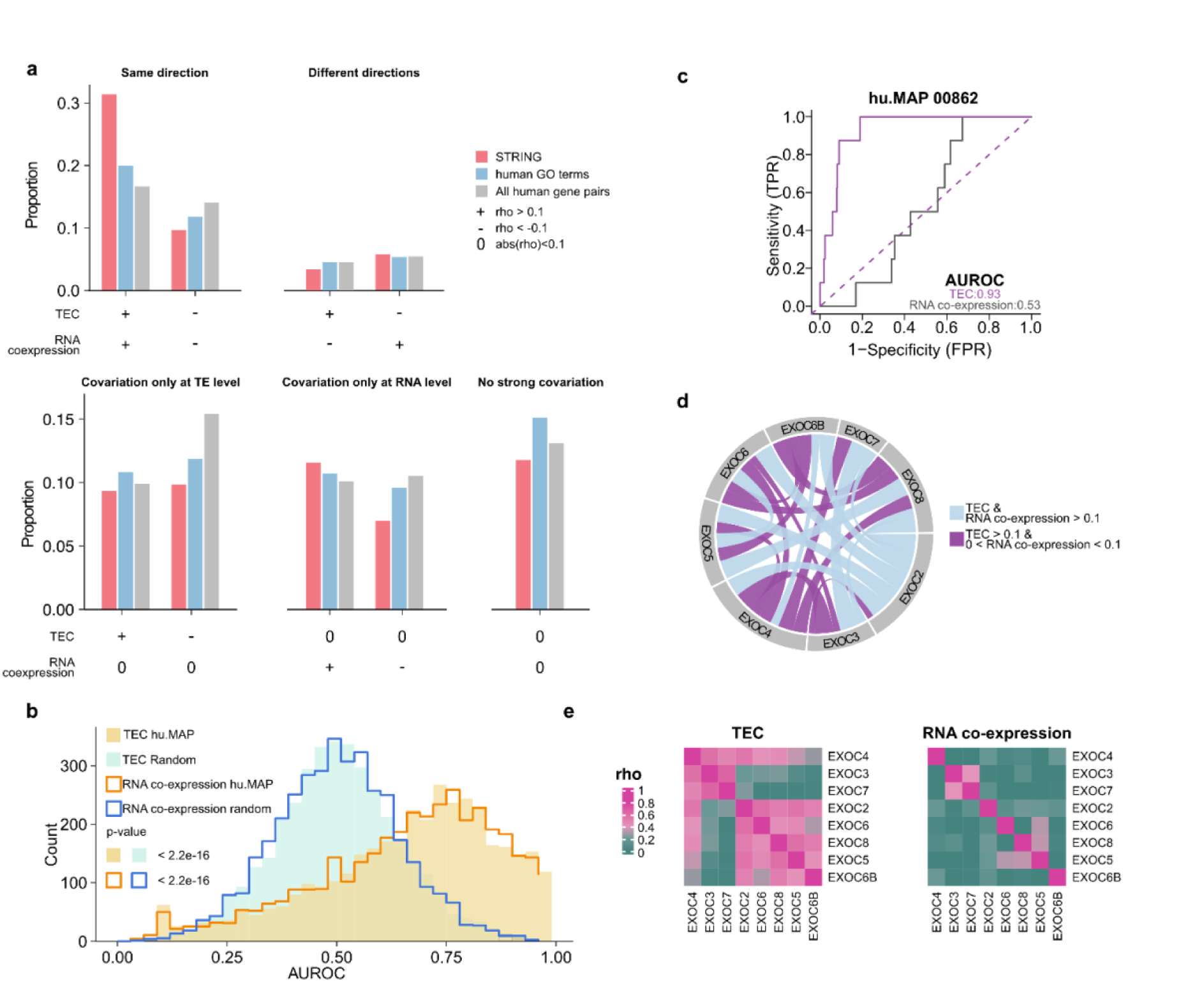
Physically interacting proteins display TEC. **a,** Number of pairs of genes among three sets (physical interaction-red; shared function-blue; all genes-gray) categorized based on the direction of similarity (Rho scores). **b,** The distribution AUROC calculated with either TEC or RNA co-expression for 3,755 hu.MAP terms (Methods). The distribution was compared to AUROC for each term that is randomly assigned genes with size matched to the original hu.MAP term. P-values were calculated using a two-sided Wilcoxon test. **c,** AUROC plot for hu.MAP term 00862, which includes eight genes within the exocyst complex. **d,** Connections represent gene pairs with rho scores above 0.1. Purple lines indicate pairs connected at TE level alone, while blue lines depict those at both the RNA co-expression and TE levels. **e,** Heatmaps display the rho calculated among genes at the TE (left) and RNA expression levels (right).

We then examined whether these patterns generalize to the higher-order organization of protein complexes. We observed protein complexes (as defined by hu.MAP^83^) displayed positive TEC and RNA co-expression (Fig. 6b; Methods). Noticeably, while proteins within the same complex generally exhibited similar positive patterns in both TEC and RNA co-expression, certain interactions within protein complexes were particularly evident only at the TE level (Fig. 6c-e).

For instance, members of the exocyst complex showed a strong positive TEC but not RNA co-expression (Fig. 6c-e). The exocyst complex consists of eight subunits in equal stoichiometry, forming two stable four-subunit modules^84,85^. Several known exocyst-binding partners are not required for its assembly and stability, indicating that the molecular details are still unclear^84^. Our finding suggests that translational regulation may play a role in maintaining the proper stoichiometry of the exocyst complex. In summary, physically interacting proteins are likely to have positive TEC in addition to positive RNA co-expression profiles. The positive correlation in RNA abundance and TE among physically interacting proteins may reflect an evolutionary pressure to efficiently utilize energy resources^24,25,86^.

## DISCUSSION

In this study, we performed a large-scale analysis of ribosome profiling and matched RNA-seq data across diverse human and mouse cell types to quantify TE. A particular challenge was inadequate metadata associated with these experiments, which limits their reuse (supplementary text). To address this issue, we conducted a manual curation process and made several methodological advances including the selection of RPF read lengths, data normalization, and estimation of TE. Although the analyzed datasets are predominantly derived from peer-reviewed publications, over 20% failed basic quality control. Our findings point to a pressing need for community standards for data quality and structured metadata.

Consistent with its established use in prior literature, we used the term “translation efficiency” to refer to ribosome occupancy per unit of mRNA (ribosome density). Recent work has questioned its interpretation as the efficiency of protein synthesis at least in the context of base-modified reporter constructs^87^. However, our work and that of others indicate TE is significantly positively correlated with protein abundance and synthesis rate for endogenous transcripts^88^. Thus, for most endogenous mRNAs, TE (ribosome density) is positively associated with protein production.

In this study, we introduced the concept of ***Translation Efficiency Covariation*** (TEC) to quantify the similarity of TE patterns across cell types. Remarkably, we found that TEC relationships are evolutionarily conserved across orthologous gene pairs in humans and mice, suggesting that translational coordination is a conserved feature of mammalian transcriptome organization. Comparative analyses of TEC and RNA co-expression networks may help distinguish shared and distinct regulatory architectures, and reveal network-level conservation and rewiring.

Importantly, TEC is predictive of gene function. Although RNA expression and TE levels are typically weakly correlated across cell types, genes with shared functions frequently exhibit coordinated patterns of both. This coordination may enhance cellular energy conservation and responsiveness to environmental cues. Analysis of TEC revealed unique insights into protein function missed by transcriptomics and proteomics. For example, LRRC28 showed strong TEC covariation with glycolytic enzymes despite absent RNA co-expression. These patterns were also undetectable at the protein-level as LRRC28 is absent from most proteomic databases such as PAXdb and ProteomeHD^44,79^.

We experimentally validated the role of LRRC28 in attenuating glycolysis in two human cell lines. Further, we identified a high confidence predicted interaction between LRRC28 and FOXK1, the key transcription factor controlling glycolytic enzyme expression, suggesting a potential mechanism. Given that LRRC28 is downregulated in multiple cancers^89–91^, its regulation may represent a previously unrecognized axis of metabolic control with therapeutic relevance ^70,92,93^.

These findings prompted us to systematically analyze TEC among physically interacting proteins, revealing a significant enrichment of positive TEC among such proteins. Coordination of TE between proteins may facilitate their co-translational assembly^81^ into complexes and contribute to their stoichiometric production, hence reducing the energetic costs of orphan protein production. These results also highlight the potential utility of TEC for synthetic biology applications. Design of synthetic gene circuits often requires precise stoichiometric production of multi-subunit complexes or pathway enzymes. In the accompanying manuscript^94^, we have developed machine learning approaches to predict TE from mRNA sequences. Such predictive modeling approaches can be combined with TEC to guide balanced translational output in engineered systems. This optimization may particularly be advantageous in applications for improving yield given that protein biosynthesis is the largest consumer of energy during cellular proliferation^25,28,86^.

We acknowledge several limitations in our study. First, the limited number of samples for certain cell lines may affect the reliability of corresponding TE estimates. Second, our analyses focused on genes robustly expressed across most cell types, limiting our ability to assess TEC for genes with cell-type-specific or consistently low expression levels. Third, due to limitations in RNA-seq quality and the short length of ribosome-protected fragments, we restricted our analysis to one representative isoform per gene^95^. As a result, we may miss regulatory features that vary across isoforms such as alternatively spliced UTRs that influence miRNAs or RBP binding. Therefore, our analyses are unable to adequately capture isoform specific usage that may affect TEC. Fourth, given the compositional nature of all sequencing experiments, TE reflects the relative allocation of translational resources. Hence, a transcript can exhibit relatively higher TE even without changes in its polysome distribution, as observed in global translational capacity shifts during viral infections and other biological contexts^96,97^. Moreover, TEC is currently defined at the level of bulk cell populations. As single-cell ribosome profiling technologies mature^98,99^, TEC could be extended to quantify translational coordination within heterogeneous cellular environments, enabling context-aware models of post-transcriptional regulation in development and disease.

In summary, TEC nominates new gene functions not captured by RNA or protein expression alone. The discovery of translation-level coordination among functionally related genes and physically interacting proteins across diverse cell types supports TEC as a conserved and functionally meaningful organizing principle of mammalian gene expression.

## ACKNOWLEDGMENTS

We thank all contributions to metadata curation: Hansel Chiang, Ashley Hoffman, Tori Tonn, Alia Segura, Charisma Tante, Eric Vasquez, and Liaoyi Xu. We also thank Yuna Shin and Victoria D. Chapman for their help with the experiments. We appreciate Dr. Milad Miladi for providing critical feedback. The original text in this paper was written by the authors. A LLM was used to suggest edits for clarity and grammar^100^. The authors acknowledge the Texas Advanced Computing Center (TACC) at The University of Texas at Austin for providing high-performance computing and storage resources that have contributed to the research results reported within this paper. URL: http://www.tacc.utexas.edu.

Research reported in this publication was supported in part by the National Institute Of General Medical Sciences of the National Institutes of Health under Award Number R35GM150667 (CC) and R35GM138340 (ESC). This work was also supported by the National Institutes of Health grant [HD110096], and the Welch Foundation grant [F-2027-20230405] (C.C.), [F-2133-20230405] (ESC). C.C. was a CPRIT Scholar in Cancer Research supported by CPRIT Grant [RR180042].

## AUTHOR CONTRIBUTIONS

Y.L., I.H., and C.C. co-wrote the original manuscript. Y.L., I.H., and S.R. generated the figures for the manuscript. H.O., M.G., and J.C. downloaded all the data from GEO and processed raw sequencing data. Y.L. and C.C. developed the translation efficiency calculation pipeline. J.C. and C.C. designed and implemented the winsorization method. Y.Z. performed the deduplication comparison. Y.L. and C.C. developed the translation efficiency covariation analysis and function prediction pipelines. Y.L., K.Q., and H.O. performed the quality control analysis for all sequencing data. Y.L. carried out covariation analysis, gene function prediction, and AlphaFold2 analysis. I.H. conducted the RBP analysis. L.P., J.W., D.Z., and V.A. assessed the quality of TE measurements by developing machine learning approaches. H.O., J.W., D.Z., V.A., Q.Z., and E.S.C. provided suggestions for the manuscript. Y.L., Q.Z., and E.S.C. conducted literature search to evaluate gene function predictions. S.R. conducted the experimental validation of LRRC28. S.R., I.H., V.G., and D.P. performed other experiments. C.C. provided study oversight, conceptualized the study and acquired funding. All authors approved the final manuscript.

## DATA AVAILABILITY

Metadata about RiboBase can be found in Supplementary table S1. Ribo files for the HeLa cell line are accessible at https://zenodo.org/records/10594392. Full TEC and RNA co-expression matrices are accessible via Zenodo repository at: https://zenodo.org/uploads/10373032. A RiboFlow configuration file and processed ribo files can be accessed at https://zenodo.org/uploads/11388478. Sequencing data and ribo files for the RBP knockout experiments are available on GEO GSE269734. Data will be publicly released upon successful review of this article.

## CODE AVAILABILITY

The code used in the study is available at https://github.com/CenikLab/TE_model/tree/main. Code will be publicly released upon successful review of this article.

## METHODS

### Acquisition and curation of ribosome profiling data

We used keyword search (“ribosome profiling”, “riboseq”, “ribo-seq”, “translation”, “ribo”, “ribosome protected footprint”) to determine studies that may employ ribosome profiling in their experimental design, from the Gene Expression Omnibus (GEO) database, with a cutoff date of January 1, 2022. Search results were manually inspected and studies containing ribosome profiling data were kept. Organism, cell line, publication, and short read archive (SRA) identifiers were obtained by automatically parsing the GEO pages of the corresponding study and sample. There was no dedicated experiment-type field for ribosome profiling experiments in GEO. Therefore we determined the experiment type (ribosome profiling, RNA-Seq, or other) of each sample by manually inspecting the GEO metadata and the associated publication of the study. Typically, ribosome profiling samples were indicated in GEO using one of the following terms: “ribosome protected footprints”, “ribo-seq”, and “ribosome profiling” in various parts of the metadata such as title, extraction protocol, and library strategy. If there were RNA-Seq samples in the same study, they were matched with ribosome profiling experiments, where available, after inspecting the sample names, metadata, and the publication of the study.

Adapters are commonly observed on the 3’ end of sequencing reads in ribosome profiling experiments, a consequence of the inherently short length of RPFs. If the 3’ adapter sequence was listed in GEO, we extracted it as part of the manual data curation process. If this sequence was unavailable, we attempted to determine it from the corresponding publication of the study. If no explicit sequence was available, we computationally analyzed the sequencing reads and searched for commonly used adapters which are CTGTAGGCACCATCAAT, AAGATCGGAAGAGCACACGTCT, AGATCGGAAGAGCACACGTCTGAACTCCAGTCAC, TGGAATTCTCGGGTGCCAAGG and AAAAAAAAAA. If any of these adapters were found in at least 50% of the reads, we used the detected sequence as the 3’ adapter. If no match was found, we removed the first 25 nucleotides of the reads anchored 6 mers and tried to extend them. If any of these extensions reached 10 nucleotides and were still detected in at least 50% of the reads, we took the highest matching sequence as the 3’ adapter. On the other hand, for sequencing reads from SRA having a length of less than 35 nucleotides, we assumed the 3’ adapters had already been removed. Detailed code can be accessed from: https://github.com/RiboBase/snakescale/blob/main/scripts/guess_adapters.py.

RiboBase was pre-populated after mining GEO. Then data curators were assigned specific studies and used the web-based interface to access the database. Each study was curated independently by at least two people. In case of disagreements, an additional experienced scientist inspected the corresponding studies and publications to make the final decision. We supplemented any missing metadata from GEO by checking the corresponding publications to ensure completeness. The result of this data curation process with information such as cell line, organism, and matched RNA-seq can be found in table S1, which forms the metadata backbone of RiboBase.

### Ribosome profiling and RNA-seq data processing

For each selected study in GEO, ribosome profiling and matching RNA-Seq reads (where available) were downloaded, from SRA, using the SRA-Tools version 2.9.1^101^, in FASTQ format using their accession numbers. FASTQ files were processed using RiboFlow^31^ where parameters were determined using the metadata in RiboBase. The reference files for human and mouse transcriptomes, annotations, and non-coding RNA sequences are available at https://github.com/RiboBase/reference_homo-sapiens and https://github.com/RiboBase/reference_mus-musculus, respectively. The detailed documentation and code used to select representative transcripts are also provided: https://github.com/ribosomeprofiling/references_for_riboflow/tree/master/transcriptome/human/v2 and https://github.com/ribosomeprofiling/references_for_riboflow/tree/master/transcriptome/mouse/v1. Briefly, the 3’ adapters of the ribosome profiling reads were trimmed using Cutadapt version 1.18^102^ and reads having lengths between 15 and 40 nucleotides were kept. Then, reads were aligned against noncoding RNAs, and unaligned reads were kept. Next, reads were aligned against transcriptome reference, and alignments having mapping quality score above 20 were kept. Reads having the same length and mapping to the same transcriptome position were collapsed, which we refer to as “PCR deduplication”. In the final step, we compiled the alignments into ribo files using RiboPy^31^. All alignment steps used bowtie2 version 2.3.4.3^103^. For each sample, we also performed the same run without the PCR deduplication step. We developed a pipeline, Snakescale, available at https://github.com/RiboBase/snakescale, to automate the entire process from downloading the data from SRA to generating the ribo files. Snakescale went over the selected list of studies and obtained their metadata from Ribobase, downloaded the sequencing data from SRA, generated Riboflow parameters file, and ran Riboflow to generate the ribo files. Examples of non- deduplicated ribo files for the HeLa cell line can be accessed at https://zenodo.org/records/10594392^104^.

To visualize the length distribution of the RPFs, we applied the scale function (z-score) in R to normalize the count of RPFs mapped to CDS regions with PCR-deduplicated ribosome profiling data. Subsequently, we plotted the distribution of these normalized RPFs using the heatmap (Fig. 1c; Supplementary Fig. 2).

### Determination of cutoff for RPF lengths and quantification of ribosome occupancy

Ribosome profiling experiments employ a range of ribonucleases including RNase I, RNase A, RNase T1, and MNase (i.e., micrococcal nuclease) (S7). These different enzymes lead to variable RPF lengths^33,105–107^. To ensure that we retain high-quality RPFs for further analyses, we implemented a dynamic extraction module that automatically selected lower and higher boundaries of RPFs for each sample. Initially, we determined the first RPF length, ranging from 21 to 40 nucleotides, that contained the highest number of CDS mapping reads. Then, we examined the two positions adjacent to this selected position. The extension of the position was carried out on either side to include a higher number of CDS-aligned reads. This extension process was repeated until it encompassed at least 85% of the total CDS reads within the 21 to 40 nucleotides range (Supplementary Fig. 3a). The final two positions identified were designated as the lower and upper boundaries. If these boundaries extended to either 21 or 40 nucleotides without including a sufficient number of reads, then 21 or 40 nucleotides, respectively, were set as the final boundaries. This approach was employed to establish the RPF cutoffs for each sample.

### Transcript coverage and quality control for ribosome profiling data

We performed quality control using RPFs that were deduplicated based on the length and position (PCR-deduplication) ribo files (Fig. 1d-e). We set two cutoffs for ribosome profiling quality control. We required that on average each nucleotide of the transcript should be covered at least 0.1 times (0.1X). Coverage was calculated with the formula:

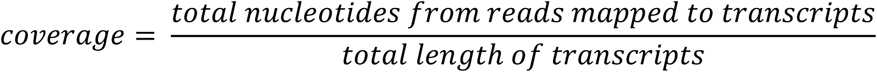

Additionally, samples with CDS mapping read percentage of 70% or higher were retained for subsequent analysis.

To assess the pattern of three nucleotide periodicity that is typically associated with ribosome profiling experiments, we first selected the length of RPFs with the highest number of counts from the PCR deduplicated ribo files. We then assigned all CDS mapping reads to one of three coding frames based on the position of their 5’ end. We aggregated the results for all genes for each sample. To facilitate comparison, we reordered the counts for each position of the three nucleotide periodicity from highest to lowest and converted these counts into percentages for each sample.

We initially classified samples based on the differences between positions 1 and 2. We identified Group 1 by selecting samples where the difference exceeded the 10th percentile of these differences between positions 1 and 2. For the remaining samples, we further classified them based on the differences between positions 2 and 3. Similarly, samples that did not exceed the 10th percentile of these differences between positions 2 and 3 among remaining samples were classified as Group 3, while the rest samples were Group 2. We further summarized the samples based on their QC status.

We classified samples from Group 1 as exhibiting three-nucleotide periodicity. The percentage of samples following three-nucleotide periodicity was calculated by dividing the number of Group 1 samples by the total number of samples across all three groups.

### PCR and Unique Molecular Identifiers (UMIs) deduplication comparison

We selected eight ribosome profiling experiments that incorporated UMIs into the sequence library preparation to assess the impact of different deduplication methods. Specifically, these samples are GSM4282032, GSM4282033, and GSM4282034 from GSE144140^108^; GSM3168387, GSM3168389, GSM3168390 from GSE115162^106^; and GSM4798525, GSM4798526 from GSE158374^96^. We processed the data using Riboflow, applying three different deduplication methods: non-deduplication, PCR deduplication, and UMI deduplication. The yaml files are available at https://github.com/CenikLab/TE_model/tree/main/riboflow_scr. The RPF length cutoffs for samples from GSE144140 and GSE115162 are listed in table S6. Since GSE158374 is not currently included in RiboBase, we manually performed the dynamic module and selected 28 to 32 as the RPF cutoff for this study.

### Winsorization of CDS mapping read counts

To address the issue of reduced usable reads resulting from PCR deduplication (supplementary text), we employed a winsorization method, which was previously proposed for tackling this problem^24,109^. For each gene’s CDS region, we obtained the distribution of non-deduplicated nucleotide counts and calculated the 99.5th percentile value. This calculation was based on reads whose lengths fell within the RPF range determined by the RPF boundary selection function. RPF counts that exceed the 99.5th percentile were capped to the value corresponding to the 99.5th percentile. This method was designed to mitigate the impact of outlier values that might arise due to disproportionate amplification during the PCR process^24^.

### Gene filtering and normalization for ribosome profiling and RNA-seq

RNA-seq experiments in RiboBase utilized several different strategies to enrich mRNAs. The two most common approaches were the depletion of ribosomal RNAs and the enrichment of transcripts by polyA-tail selection. This difference leads to dramatically different quantification of a subset of genes that lack polyA-tails (e.g. histone genes, Supplementary Fig. 6c). Hence, we removed 166 human and 51 mouse genes identified as lacking polyA tails (table S8-9)^110,111^.

We normalized both PCR-deduplicated RNA-seq data and winsorized non-deduplicated ribosome profiling data with counts per million (CPM) after removing the genes without polyA-tails. Genes with CPM less than one in over 30% of the total samples in both RNA-seq and ribosome profiling for either human or mouse were removed in further analyses. 11,149 human and 11,434 mouse genes were retained using this approach. We have summed the counts of all polyA genes that were filtered out and grouped them under ‘others’ in the count table.

### Validation of manual curation and quality control by matching between RNA-seq and ribosome profiling from RiboBase

We assessed the manual matching of ribosome profiling (winsorization) and RNA-seq (PCR deduplication) data in RiboBase by establishing a matching score for the samples that successfully passed quality control (transcript coverage > 0.1X and CDS percentage > 70% with PCR-deduplicated ribosome profiling data). We calculated the coefficient of determination (R²) using the Centered Log Ratio (CLR) transformed gene counts. This was done for each ribosome profiling sample against all corresponding RNA-seq samples within the same study. Subsequently, for each ribosome profiling sample, we calculated the difference between the R² of its matching pair from RiboBase and the mean R² of the non-matching pairs within the same study. The difference was defined as the matching score.

To remove poorly matched samples in both human and mouse datasets, we established a cutoff based on the R² from the matched ribosome profiling and RNA-seq data in RiboBase. Any sample with an R² lower than 0.188 in either human or mouse, which is Q1 - 1.5 * IQR of mouse R² distribution, was considered a poor match and consequently excluded from further analysis (Supplementary Fig. 6b). Finally 1,054 human and 835 mouse ribosome profiling experiments with their matched RNA-seq were used for TE calculation.

### Translation Efficiency (TE) calculation

CLR normalized counts from PCR-deduplicated RNA-seq and winsorized non-deduplicated ribosome profiling were used to calculate TE with compositional linear regression^36,112,113^. Missing values can limit the power of TE calculations, and there is ongoing debate on how to address zeros in compositional data ^114^. Previous studies have shown that approximately 15–40% of genes in bulk RNA-seq datasets from various tissues are not expressed ^115^. To address the issue of missing values, we followed the guidance from the propr library, replacing zeros with the lowest observed read count (one). This approach ensures that these zeros are represented as a low proportion within the sample. We also applied two other zero imputation methods: Geometric Bayesian-Multiplicative and Square-root Bayesian-Multiplicative from zCompositions package (version:1.5.0.4) for TE and TEC calculations.

In our linear regression approach, ribosome profiling data served as the dependent variable, while the corresponding RNA-seq data provided the explanatory variable. The first step involved transforming the gene count, which includes ‘others’, into CLR normalized compositional vectors. Given the constraints of count data within a simplex, a further transformation from CLR to Isometric Log Ratio (ILR) was necessary for linear regression^36^. This transformation is crucial as it allows the compositional data to be decomposed into an array of uncorrelated variables while preserving relative proportions. The ILR transformation projects the original data onto a set of orthonormal basis vectors derived from the Aitchison simplex. Then the linear regression model applied to these transformed variables can be represented as:

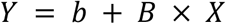

Where Y is the ILR-transformed ribosome profiling data and X is the ILR-transformed RNA-seq data. The model assumes a normal distribution:

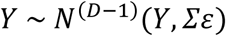

Where Σε represents the residual variances. These residuals were then extracted from each sample and reconverted to CLR coordinates which are used as the definition of TE for each gene in each sample. Finally, we averaged TE for different cell lines and tissues (Fig. 2b, Supplementary Fig. 7), and reported the TE in table S10-11. The scripts to generate TE are available at https://github.com/CenikLab/TE_model. We also provide uncertainty estimates calculated using the metric standard deviation designed for compositional data from the Compositions (version: 2.0.8) library (table S21-22).

### Correlation between translation efficiency and protein abundance

We assessed the correlation between TE and protein abundance from seven human cell lines (A549, HEK293, HeLa, HepG2, K562, MCF7, and U2OS). The protein measurements were obtained from PAXdb^44^. 9924 genes were shared between our TE and the protein abundance data. We calculated the Spearman correlation coefficient for each cell line using the ‘stats’ package in R to evaluate the relationship between TE and protein abundance. For a more detailed discussion of how to interpret similarity between protein abundance and sequencing based measurements of RNA-expression and translation see Csardi et al.^116^.

### Conservation of translation efficiency between orthologous genes from human and mouse

Orthologous genes between human and mouse were identified using the ‘orthogene’ package from Bioconductor^117^ using the parameters ‘standardise_genes=TRUE, method_all_genes=“homologene”, non121_strategy=“keep_both_species”’. A single human gene could correspond to multiple mouse orthologs or vice versa. To maintain all one-to-many matches in our analysis, each correspondence is represented by multiple rows in our table (if a human gene ‘A’ is orthologous to mouse genes ‘B’ and ‘C’, we generate two separate rows: ‘A-B’ and ‘A-C’). Human genes lacking corresponding mouse orthologs were excluded or vice versa. As a result, a total of 9,194 gene pairs were identified as orthologous between human and mouse (table S12)

To capture the variability in TE and mRNA expression between orthologous genes in human and mouse, we measured the standard deviation using the metric standard deviation (msd) function from the ‘compositions’ package in R^118^. We observed a negative Spearman correlation coefficient between msd of TE and mean TE, as well as msd of RNA expression and mean RNA expression, in both species. To address the dependency between msd and mean values, we conducted a partial correlation analysis. For example, we adjusted the human msd values using the mean TE from both human and mouse with the ‘pcor.test’ function from the ‘ppcor’ package^119^.

GO term analysis was performed using FuncAssociate 3.0, accessible at http://llama.mshri.on.ca/funcassociate/^120^. For this analysis, we set either 9,194 mouse or 9,189 human orthologous genes as the background. We generated association files for these genes with the December 4, 2022 version of human or mouse GO terms. In the human or mouse association file, we only kept those GO terms containing at least 10 genes for further analysis.

### Assessment of methods for calculating genes’ similarity with ribosome occupancy data

We used eight commonly used methods to quantify the similarity of ribosome occupancy across cell types for all pairs of 11,149 human or 11,434 mouse genes in RiboBase.

Method 1 - CPM-normalized ribosome footprint counts were used to calculate the Pearson correlation coefficient as implemented in the stats R package.

Method 2 - Quantile-normalized (customized Python script) ribosome footprint counts were used to calculate the Pearson correlation coefficient.

Method 3 - Ranking of ribosome footprint counts was used to calculate the Spearman correlation coefficient as implemented in the stats R package.

Method 4 - CLR-normalized ribosome footprint counts were used to calculate the proportionality (rho scores) between genes as implemented in the propr package with lr2rho function^38^.

Method 5 - CPM-normalized ribosome footprint counts were used to calculate the similarity between genes with a decision tree-based method as implemented in the treeClust package^79,121^. We applied the ‘treeClust.dist’ function with a dissimilarity specifier set to d.num=2.

Method 6 - Quantile-normalized ribosome footprint counts were used to calculate the similarity between genes with the decision tree-based method.

Method 7 - CPM-normalized ribosome footprint counts were used to calculate gene similarity with the generalized least squares (GLS) method^122^.

Method 8 - Quantile-normalized ribosome footprint counts were used to calculate gene similarity with the GLS method.

We compared these eight ribosome occupancy similarity matrices to determine the most effective method for constructing gene relationships with respect to biological functions. This assessment employed the guilt by association principle to ascertain the functional coherence within a gene matrix, determining if genes associated with a particular biological function (GO terms^123^, TOP mRNAs^124^) exhibit similar expression patterns and network interactions^125^.

The complete ontology was sourced from the Gene Ontology website, with the files goa_human.gpad.gz and mgi.gpad.gz, generated on December 4, 2022^123^. The annotation of Gene Ontology terms was accomplished with the aid of the org.Hs.eg.db and org.Mm.eg.db R packages^126,127^. We restricted the selection of GO terms to those associated with the 11,149 human and 11,434 mouse genes that had passed gene filtering. We used GO terms associated with at least 10 but less than 1,000 genes for evaluation, yielding a total of 2,989 human and 3,340 mouse GO terms.

We then employed the neighbor-voting algorithm to assess the covariations of ribosome occupancy among genes from the same GO term with AUROC^125^. Specifically, we first converted the similarity scores to absolute values. Then we extracted genes associated with a specific function and implemented the leave-one-out cross-validation method. For this analysis, we iteratively masked one gene at a time, treating it as if it did not belong to the function. In each iteration, we calculated the total sum of similarity scores from all genes not belonging to the function to all the remaining genes within the function. We normalized the sum of similarity scores for each gene against the sum of similarity scores for that gene with all genes. After normalization, we converted these normalized similarity scores into rankings. We retained the rankings only for genes that belong to this specified functional property. Finally, we computed the AUROC for all genes within this functional property based on these rankings. A detailed script for genes’ functional similarity pattern analysis can be found: https://github.com/CenikLab/TE_model/blob/main/other_scr/benchmarking.R.

### RNA co-expression and translation efficiency covariation

We introduce the concept of TEC, which employs a compositional data analysis approach^36,38^ to quantify the similarity patterns of TE across various cell and tissue sources, as described in Method 4 above. The proportionality scores were calculated with the following formula from the propr package with lr2rho function^38^:

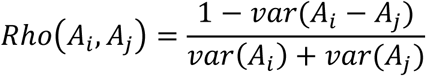

Where Ai and Aj represent TE values for genes i and j from the TE matrix A.

In this study, the TEC was calculated with 77 human cell lines for 11,149 genes or 68 mouse cell lines for 11,434 genes. The proportionality coefficients (rho scores) generated from this method range from -1 to 1. Full TEC and RNA co-expression matrices are accessible via Zenodo repository at: https://zenodo.org/uploads/10373032.

### Evaluation of the ability of TEC to predict novel gene functions

We compared the AUROC between an older version of GO terms (January 1, 2021) to the newer version of GO terms (December 4, 2022) to identify genes that had been newly added to the GO terms in this timeframe. GO terms were downloaded and filtered to include only those terms containing between 10 and 1,000 genes with either human or mouse backgrounds (11,149 human genes or 11,434 mouse genes). We selected 184 human and 238 mouse GO terms from the older version that demonstrated high TEC similarity (AUROC > 0.8) among genes within the same term for predicting novel gene functions. We first converted the rho scores for TEC between gene pairs to absolute values. For genes not currently included in the GO terms, we calculated the sum of rho for each gene relative to all genes within the term, based on either TE or mRNA expression levels. We then normalized these rho sums for each gene against the total rho sum of that gene across all 11,149 human genes or 11,434 mouse genes. These normalized values were converted into ranking percentages to reflect the likelihood of these genes being associated with the respective GO term. Finally, we identified the top-ranking genes as potentially new additions and cross-validated them with the newer version of the GO terms to confirm our predictions.

### Prediction of novel gene functions with TEC

We analyzed 243 human and 310 mouse GO terms as of December 4, 2022, which demonstrated high similarity patterns between genes in TE level (AUROC > 0.8) to predict novel gene functions. Absolute TEC rho scores served as the input for biological function prediction (GO terms). The prediction method followed the same protocol as our previous evaluations of TEC’s ability to predict novel gene functions. However, we added a filter step: a newly predicted gene was retained only if its average rho score with other genes within the same term exceeded the overall average rho score for all existing genes in that term. This prediction analysis was performed using a custom script that can be found at https://github.com/CenikLab/TE_model/blob/main/other_scr/prediction.R.

### Computational evaluation of the interaction between LRRC28, glycolytic proteins, and proteins from forkhead TF family

We computed the pair-wise interaction probabilities between LRRC28 or LRRC42 and glycolytic proteins (HK1, HK2, PFKL, PFKM, PFKP, TPI1, PGK1, ENO1, ENO2, and PKM) with AlphaFold2-Multimer 2.3.0^72,128^. In addition, we also calculated pairwise interaction probabilities for LRRC28 with 35 proteins from the forkhead transcription TF family^78^. We extracted the canonical amino acid sequence for each gene from UniPort^129^ as the input file. We set 0.7 as the cutoff of ipTM+pTM as a high-confidence protein structure and binding probability cutoff^75^. We then evaluated the interfaces predicted by AlphaFold2-Multimer, using a pDOCKQ score greater than 0.23 as our criterion for reliability^76,77^.

### Benchmarking TEC and RNA co-expression for protein interactions

Using a similar approach to our benchmarking with biological functions, we employed the neighbor-voting algorithm to assess physical protein interactions based on rho scores among genes at either the TE or mRNA expression level. We first kept the non-negative rho between genes and set negative rho to zero. We then analyzed similarity patterns between genes from the same protein complex, downloading from the hu.MAP 2.0 website^83^. In this process, we excluded genes from hu.MAP terms that were not in the 11,149 human gene list, resulting in 8,024 overlapping genes between our list and hu.MAP terms. Furthermore, we removed hu.MAP terms that included fewer than three genes. This filtering process left us with 3,880 hu.MAP terms, among which 3,755 contained unique genes.

Since proteins within the same complex may not physically interact, we used physical interaction pairs downloaded from the STRING website instead of gene pairs from hu.MAP terms to summarize the interactions in Fig. 6a.

### Identification of enriched RNA motifs among genes with high degree of TEC

To reduce bias in motif enrichment analysis that may arise by ribosome footprint mapping to paralogous genes, we removed predicted paralogs from each GO term using Paralog Explorer^130^ (DIOPT score > 1). Then, we enumerated heptamers in each transcript region using the Transite kmer-TSMA method^60^ with default parameters for each species (human, mouse), transcript region (5’ UTR, CDS, 3’ UTR), and GO term (selected terms with TE AUROC > 0.7, TE-RNA AUROC difference > 0.2, and number genes after paralog removal >= 12). We selected the three mouse terms and top five terms in humans with the highest number of genes and greatest AUROC difference.

After counting heptamers with Transite, we selected motifs that had >20 hits among genes in the GO term to address assumptions of uniformity near p-values of 1 for some multiple-test correction methods. Then, we used the Holm method to correct p-values for each species separately, and selected motifs with an adjusted p-value < 0.05. Finally, heptamers were annotated with RBPs included in the Transite^60^ and oRNAment databases^59^. For annotation of RBPs in the oRNAment database, we required that the heptamer have a matrix similarity score^59^ of 0.8 or greater when matching to each RBP position weight matrix. RBP motif hits from other species (*Drosophila*, artificial constructs) were removed from RBP annotations, and the hits to the heptamer of *Drosophila* tra2 were annotated as TRA2A for human genes with the term GO: 0140678.

eCLIP data for PABPN1, SRSF1, and TRA2A were downloaded from ENCODE^61^ as BED files (K562 and HepG2 cell lines, GRCh38 reference). The BED files for biological replicates were concatenated and peaks that overlapped by at least one base pair were merged with ‘bedtools merge -s -c 4,6,7 -o collapse’^131^. The resulting merged peaks were intersected with transcripts in the GO term of interest and an equal number of control transcripts (Gencode v34 GTF). The control transcripts were selected by matching on length and GC content for each transcript region (5’ UTR, CDS, 3’ UTR) using MatchIt^132^ with default parameters. Because the gene *CARMIL2* in GO term GO:0010592 does not have a 5’ UTR, required for matching, we assigned it a dummy 5’ UTR with length and GC content equal to the median across all transcripts. The number of eCLIP peaks in the CDS for each RBP were summed for genes in the GO term and control genes.

### Identification of RBP-transcript pairs with high correlation between RBP RNA expression and transcript TE

The Pearson correlation coefficient between transcript TE and the RNA expression of RBPs from human and mouse^51^ was tested using R stats::cor.test after taking the mean of these values by cell types and tissues. P-values were corrected with the Benjamini-Hochberg procedure, and correlations were deemed significant at a FDR < 0.05. For cloning the guides required for knockout cell line generation, top two ranked guides were selected from the Brunello library^133^ for each RBPs (table S19). The guides were cloned in LentiCRISPRv2 (Addgene, 52961) as per protocol^134^ and confirmed by Sanger sequencing. Briefly, for lentiviral production, HEK293T cells were seeded at a density of 1.2 x 10^6^ cells per well in a 6-well plate in OPTI-MEM media supplemented with 5% FBS and 100 mM Sodium Pyruvate, 24 h prior to transfection. Both the cloned gRNA plasmids for each RBP (700 ng of each transfer plasmid) were co-transfected with the packaging plasmids pMD2.G and psPAX2 (Addgene; 12259 and 12260) using Lipofectamine 3000 (Invitrogen) and the virus was collected as per the manufacturer’s protocol. For generation of the knockout clones, HEK293T cells were seeded at a density of 5 x 10^4^ cells per well in a 6-well plate in DMEM media supplemented with 10% FBS, 24 h prior to infection. Next day, the media was replaced with 1.5 ml of 1:2 diluted lentivirus containing polybrene (8 µg/mL). After 16 h, the lentivirus was replaced with fresh media and puromycin (2 µg/mL) was added to the cells 48 h after transduction. The selection continued for 5 days followed by a period of recovery for 24 h before harvesting the cells.

### Ribosome profiling and RNA sequencing of RBP knockout cell lines

Three million cells for the PARK7, USP42, and VIM knockout cell lines, along with an AAVS1 (safe harbor control) knockout line, were plated in three 10 cm^2^ dishes. 27 h later cells at ∼60% confluency were treated with 100 µg/mL cycloheximide (CHX) for 10 min at 37 °C, then collected in ice-cold PBS with 100 µg/mL CHX. Cells were spun at 100 x g for 7 min at 4 °C, then flash frozen in liquid nitrogen and stored at -80 °C. Cell pellets were lysed with 400 µL lysis buffer (20 mM Tris-HCl pH 7.4, 150 mM NaCl, 5 mM MgCl2, 1 mM DTT, 1% Triton-X, 100 µg/mL CHX, 1x protease inhibitor EDTA free) for 10 min on ice. Lysates were clarified by centrifugation at 1,300 rpm for 10 min at 4 °C. 40 µL lysate was saved for total RNA extraction, and the rest of the lysate was digested with 7 µL RNaseI for 1 h at 4 °C. Digestion was stopped by adding ribonucleoside vanadyl complex to a final concentration of 20 mM. Digested lysates were then loaded onto a 10 mL sucrose cushion (20 mM Tris-HCl pH 7.4, 150 mM NaCl, 5 mM MgCl2, 1 mM DTT, 1 M sucrose) and centrifuged at 38,000 rpm for 2.5 h at 4 °C using a SW41-Ti rotor. The pellet and the total RNA aliquot were both solubilized with 1 mL Trizol, and RNA was purified with the Zymo Direct-zol RNA Miniprep Kit, including DNaseI digestion.

Quality of total RNA was confirmed with Bioanalyzer RNA Pico. All RIN scores were >= 9.8. Libraries were prepared from 1 µg total RNA using the NEBNext Ultra II RNA Library Prep Kit for Illumina according to manufacturer’s protocol and using 8 cycles for PCR.

RPFs were size-selected on a 15% TBE urea gel by electrophoresing at 150 V for 1.5 h. RPFs between 28-32 nt were sliced, using 28 nt and 35 nt markers as a guide. Slices were frozen at -20 °C for 1 h, crushed with pestles, and the RPFs were eluted in gel extraction buffer (300 mM sodium acetate pH 5.5, 5 mM MgCl2) by rotating overnight at room temperature. Eluates were passed through Costar Spin-X filter tubes at 12,000 x g for 1 min 30 s. Then 1 µL 1 M MgCl2, 2.5 µL GlycoBlue, and 1 mL ethanol were added and the RPFs precipitated for two days at -20 °C. Pellets were dried and resuspended in 16 µL water.

Libraries were generated from 8 µL RPF eluate using the Diagenode D-Plex Small RNA Kit with minor modifications: in the 3’ dephosphorylation step 0.5 µL T4 PNK was supplemented and incubated for 25 min. The RTPM reverse transcription primer was used and 8 cycles were performed for PCR. Libraries were quantified by Bioanalyzer High Sensitivity DNA Kit, pooled equimolar according to the quantity of the peak for libraries with full-length inserts (∼204 nt), and cleaned up with 1.8X AMPure XP beads. Adapter dimers and empty libraries were removed by size selection on a 12% TBE PAGE gel, followed by extraction with the crush and soak method, and final libraries were resuspended in 20 µL water.

### Ribosome profiling and RNA sequencing analysis for RBP knockouts

Analysis was conducted using RiboFlow v0.0.1 with deduplication of both Ribo-seq and RNA-seq data. A RiboFlow configuration file and processed ribo files can be accessed at https://zenodo.org/uploads/11388478.

We used edgeR to measure RBP KO effects on 1) RNA abundance and 2) gene TE. To do this, we respectively modeled 1) RNA-seq counts of a specific RBP KO line to that of the other two RBP KO lines; and 2) Ribo-seq counts, contrasted with RNA-seq counts, for a specific RBP KO line compared to the other two RBP KO lines. All counts were enumerated from mapped reads to the coding regions. We originally included a control KO line (AAVS1 locus) for comparison; however, by PCA, this KO line showed a distinct gene expression signature from that of the other KO lines, indicating it may not be suitable as a control (Supplementary Fig. 16d-e). Further, using the AAVS1 KO line as a control, we observed highly similar hits for each RBP KO tested. We included filtering of counts using edgeR::filterByExpr with default parameters, the TMM method for calculation of size factors, and quasi-likelihood negative binomial models for fitting. Genes were considered differential at FDR < 0.05.

### Cell Culture

HEK293T and MCF-7 cell lines were obtained from ATCC. Human hepatoma Huh-7.5 cells were a gift from Charles Rice, Rockefeller University and human SH-SY5Y neuroblastoma cells were obtained from Tanya Paull (University of Texas at Austin, Austin, TX, USA). All cell lines were maintained in Dulbecco’s modified Eagle’s media (DMEM, GIBCO) supplemented with 10% Fetal Bovine Serum (FBS, GIBCO, Life Technologies) and 1% Penicillin and Streptomycin (GIBCO, Life Technologies) at 37°C in 5% CO2 atmosphere. Cell lines were tested for mycoplasma contamination every 6 months.

### Plasmid cloning

pCMV SPORT6 vector with LRRC28 was obtained from transOMIC (TCH1003). LRRC42 ORF was amplified using cDNA from HEK293T cells. The oligos used for PCR amplification of the two genes had a FLAG sequence to be incorporated as a N-terminal tag along with BamHI and EcoRI restriction sites in forward and reverse primers, respectively.

PCR amplification of the LRRC28 and LRRC42 ORF using the above mentioned templates were carried out using Q5 polymerase (NEB) with the following protocol; 98 °C for 30 sec, 30 cycles of 98 °C for 10 sec, 55 °C for 10 sec and 72 °C for 40 sec and a final extension at 72 °C for 2 min. The PCR amplified products were gel extracted followed by double digestion of the amplicon and the pLVX-M vector (Addgene, 125839) with EcoRI -HF (NEB) and BamHI-HF (NEB) at 37 °C for 1 h. Post digestion, the fragments were purified and cloned using T4 DNA ligase at a ratio of 1:3 at 25 °C for 1 h followed by heat inactivation at 65 °C for 15 min and transformation by heat shock method. For LRRC42 cloning, ORF was amplified using cDNA from HEK293T cell line and two sequences (seq1 and seq 2) from the pCMV-SPORT6 adjacent to EcoRI and NotI site were amplified with LRRC42 overhangs using oligos in table S24. The vector; pCMV SPORT6 vector with LRRC28 was digested with EcoRI and NotI followed by cloning of LRRC42 and two sequences by Gibson cloning. All clones were verified by Sanger sequencing (ACGT Inc.) and full plasmid sequencing (Plasmidsaurus).

### Lentivirus and stable cell line generation

For lentiviral generation, 1.2 X 10^6 HEK293T cells were seeded in a 6-well plate followed by transfection of the cloned plasmid along with psPAX2 (Addgene, 12260) and pMD2.G (Addgene, 12259) plasmids using Lipofectamine 3000 (Invitrogen) as per the manufacturer’s protocol. The generated lentivirus was used for infection using 8 μg/ ml of polybrene for HEK293T, MCF-7 and Huh 7.5 and 2 μg/ ml of polybrene for SH-SY5Y. After 48 h of infection, puromycin was added at concentrations of 2μg/ ml for HEK293T, Huh 7.5 and 1μg/ ml for MCF-7, SH-SY5Y. The selection was continued for 5 days following recovery in antibiotic-free media.

### Measurement of extracellular acidification rates (ECAR)

20,000 cells of HEK293T, MCF-7 and 40,000 cells of SH-SY5Y stably expressing LRRC28 or LRRC48 and 20,000 cells of Huh 7.5 were seeded onto poly-L lysine coated XF Pro Cell Culture Microplates with Dulbecco’s modified Eagle’s media (DMEM, GIBCO) supplemented with 10% Fetal Bovine Serum (FBS, GIBCO, Life Technologies). Huh 7.5 was transiently transfected with either pCMV SPORT6 vector with LRRC28 or LRRC42 using lipofectamine 3000 as per manufacturer’s protocol. The ECAR was measured using Seahorse XF Pro (Agilent). The cartridges were hydrated a day prior to the experiment as per the manufacturer’s instructions. On the day of the experiment, prior to performing the assay, the cells were washed with Seahorse XF DMEM Medium pH 7.4 (Agilient, 103575), supplemented with 2 mM of L- Glutamine (Agilient; 103579-100) followed by incubation in non-CO2 incubator for 1 h. To determine glycolytic capacity of cells expressing LRRC28 or LRRC42, glucose (Agilient, 103577-100) was injected at a final concentration of 10 mM (Injection 1) followed by injection of 100 mM 2-Deoxy-D- Glucose (Sigma, D6134-1G) (Injection 2). Three measurement cycles for each assay point i.e. baseline, injection 1 and 2 were carried out with each measurement cycle consisting of a mixing time of 3 minutes and a data acquisition period of 3 minutes. The results were analyzed using Wave Pro 10.2.1.

