## Extended Figures for "Translation efficiency covariation across cell types is a conserved organizing principle of mammalian transcriptomes"

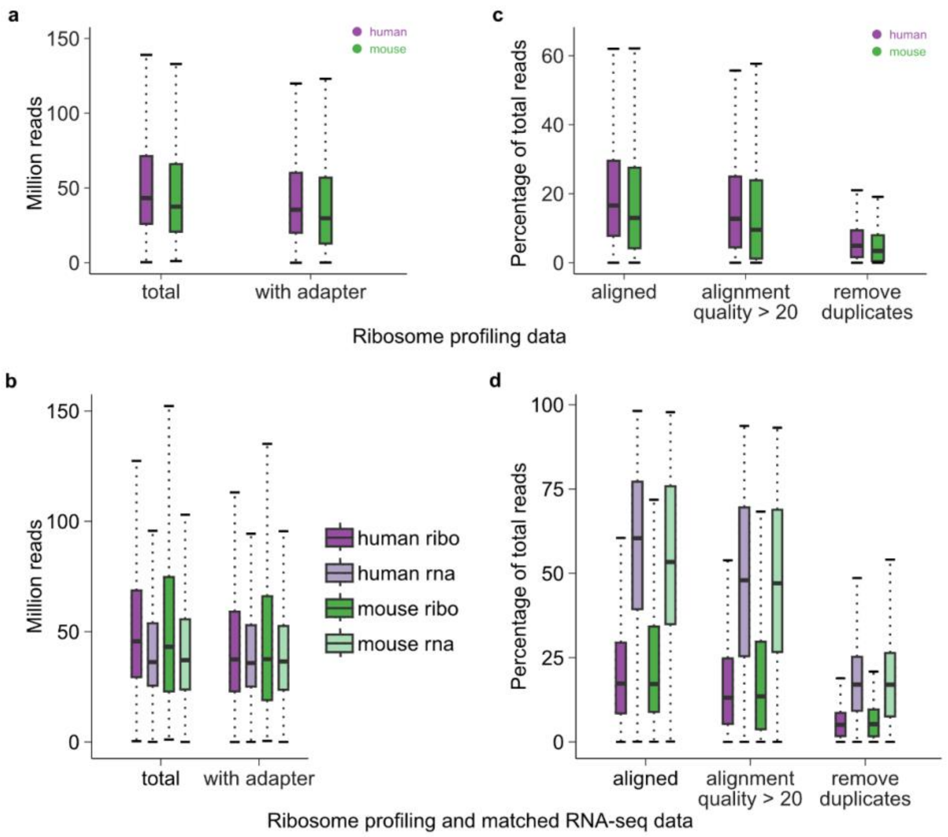


**Supplementary Fig. 1 | Sequencing quality of ribosome profiling data with matched RNA-seq data when available (supplementary text): a,** Distribution of read counts for ribosome profiling data in RiboBase. In all figure panels, the horizontal line corresponds to the median. The box represents the interquartile range and the whiskers extend to 1.5 times of it. **b,** Distribution plot similar to panel A for ribosome profiling data with matched RNA-seq. **c,** Distribution of the proportion of read count aligned to transcripts, read counts with high-quality alignments, and the percentage of reads remaining after PCR deduplication, relative to the total number of reads from panel A. **d,** Similar plot as panel C for ribosome profiling with matched RNA-seq.

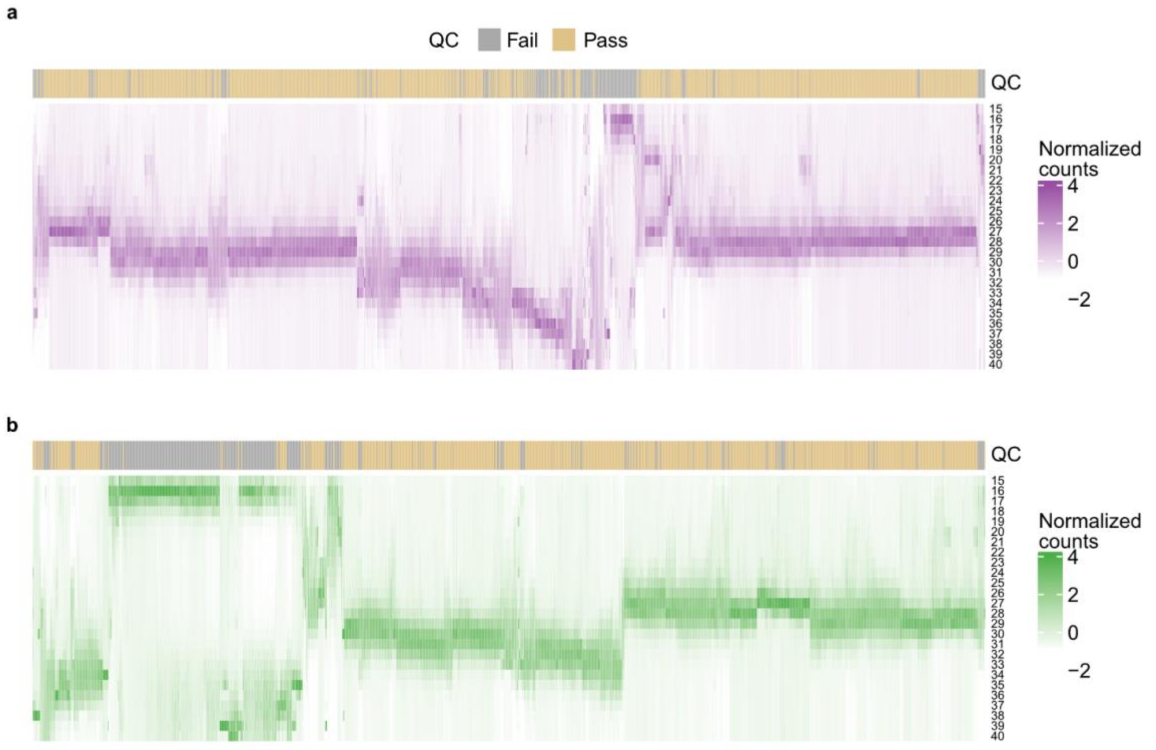


**Supplementary Fig. 2 | Length distribution of RPFs for human and mouse samples: a,** The read length distribution of RPFs aligned to coding sequences for all human experiments. The color in the heatmap represents the z-score adjusted RPF counts (Methods). Each experiment where the percentage of RPFs mapping to CDS was greater than 70% and achieving sufficient coverage of the transcript (>= 0.1X) was annotated as QC-pass. **b,** Similar to panel A for mouse samples.

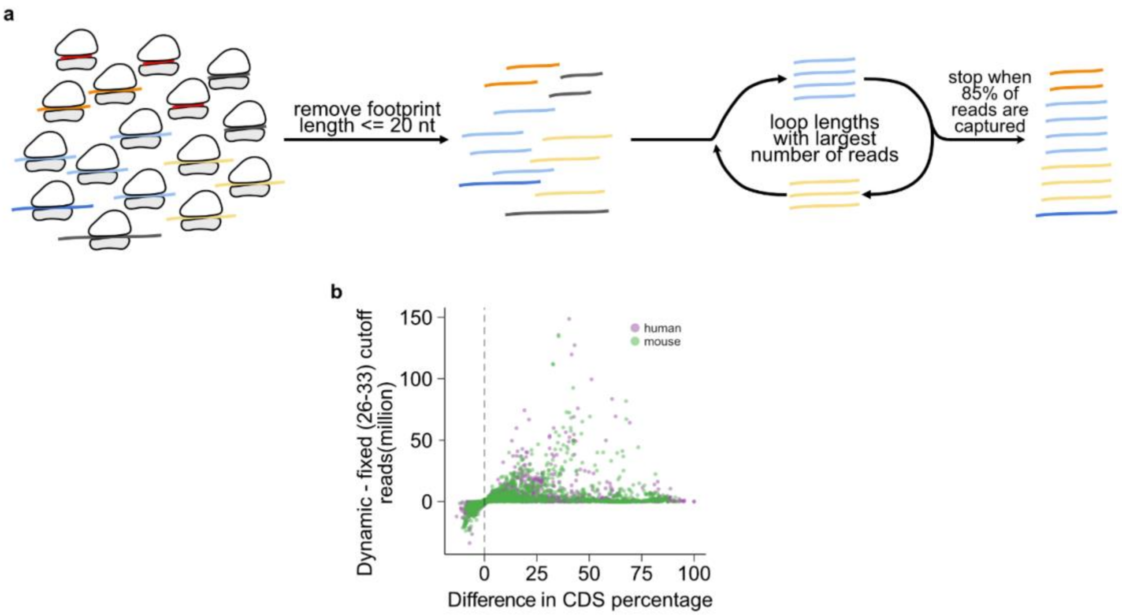


**Supplementary Fig. 3 | Schematic for method to select range of RPF lengths: a,** RPFs shorter than 21 nucleotides were removed, then we identified the RPF length with the highest number of reads mapping to CDS to serve as the starting point. Subsequently, we compared one nucleotide longer or shorter than the first and chose the length with the most reads again. This looping process continued until at least 85% of the total CDS mapping RPFs were included. **b,** We compared the usable reads selected with two different boundary cutoffs (y-axis) and the proportion of these selected reads that map to the coding regions (x-axis) for each ribosome profiling experiment.

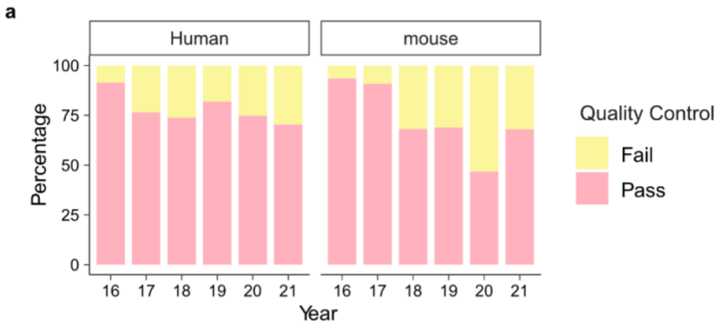


**Supplementary Fig. 4 | Data quality of ribosome profiling experiments from 2016 to 2021: a,** The percentage of ribosome profiling experiments from GEO that pass or fail quality control (the percentage of RPFs mapping to CDS was greater than 70% and achieving at least 0.1X coverage of the transcript as QC pass).

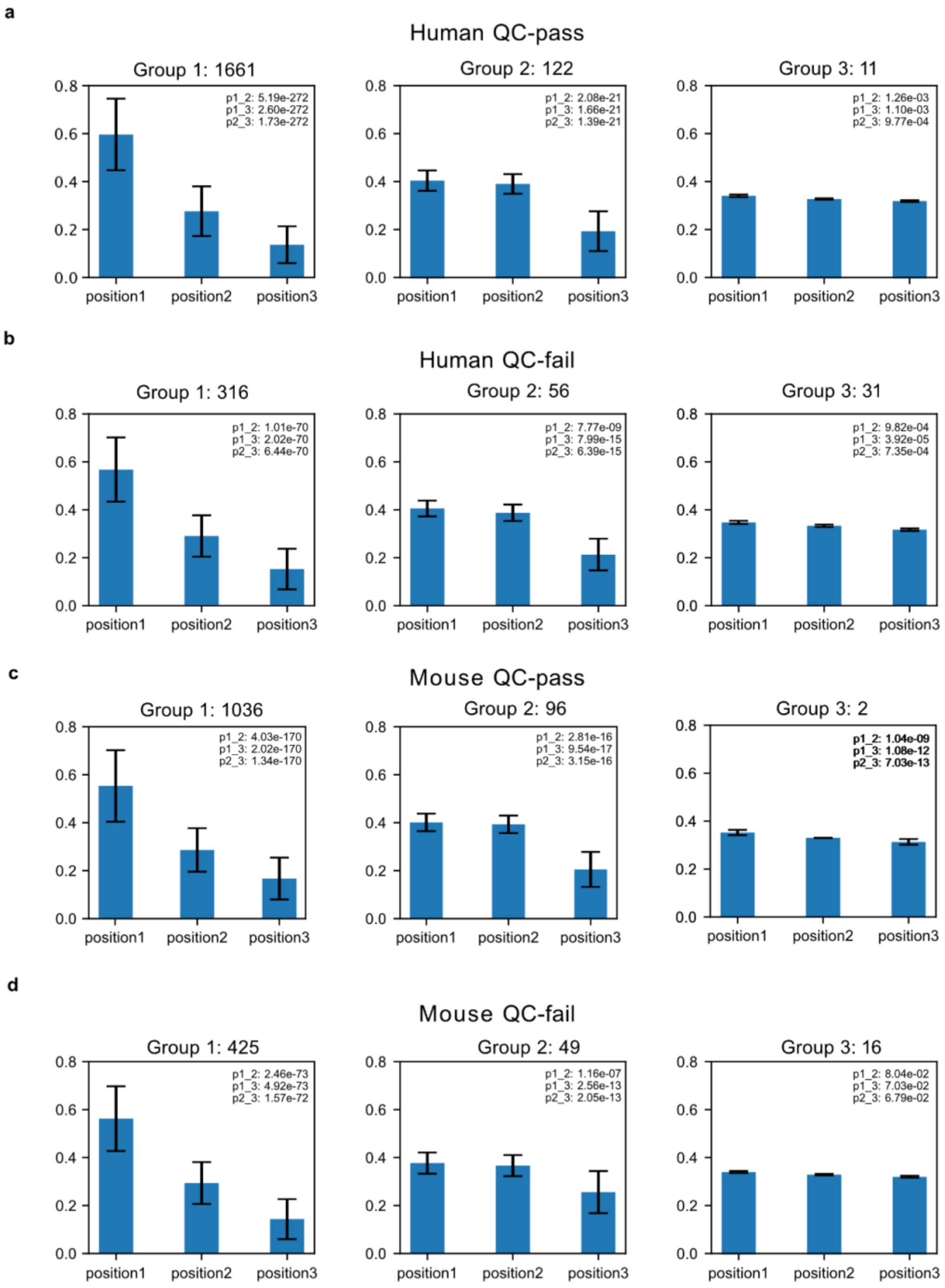


**Supplementary Fig. 5 | Three nucleotide periodicity of ribosome profiling data: a-d,** In ribosome profiling experiments from RiboBase, samples were classified according to distinct periodicity patterns (Methods). For all figure panels, we added error bars to represent the standard deviation across samples. Statistical significance was assessed using the Wilcoxon test, and the p-values were subsequently adjusted for all 33 comparisons using the Benjamini-Hochberg method. We considered the Group 1 pattern as indicative of the expected three-nucleotide periodicity patterns.

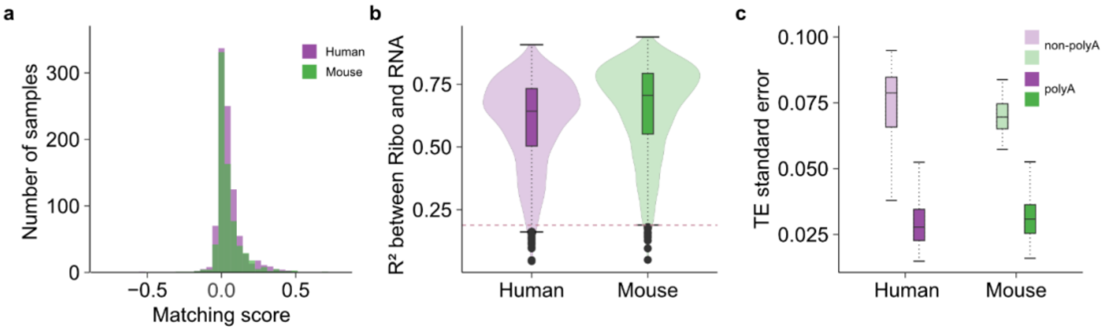


**Supplementary Fig. 6 | Validation of ribosome profiling and RNA-seq matching and gene selection for TE calculation: a,** We calculated the coefficient of determination ($R^2$) between a specific ribosome profiling experiment and its corresponding RNA-seq from RiboBase. Additionally, we determined the average $R^2$ for all other pairings for the same ribosome profiling sample with other RNA-seq data from the same study. The matching score represents the difference in $R^2$ values between these two (x-axis; Methods). **b,** A dashed line at 0.188 serves as the threshold to identify samples with poor matching. In each figure panel containing boxplots, the horizontal line corresponds to the median. The box represents the IQR and the whiskers extend to 1.5 times of it. **c,** Distribution of standard error of TE values across tissue and cell lines (y-axis) for genes with polyA and without polyA tails.

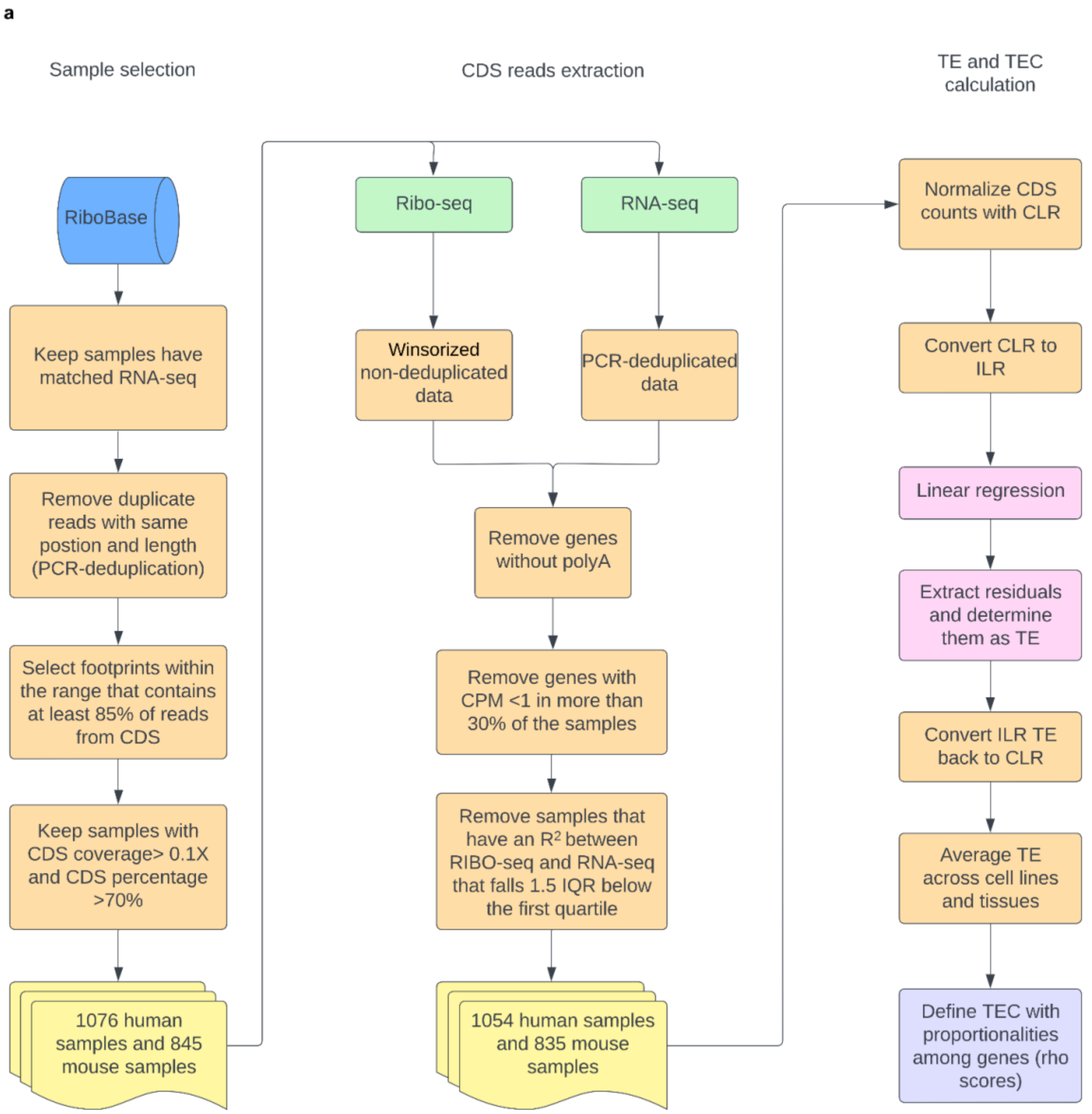


**Supplementary Fig. 7 | Detailed workflow of data processing for TE and TEC calculations:** **a,** We selected ribosome profiling data with matched RNA-seq and removed duplicated reads with identical positions and lengths (PCR-deduplication). We set the RPF read length range for individual samples with our dynamic cutoff and filtered out ribosome profiling experiments that failed quality control. After selecting high-quality samples, we reprocessed all these ribosome profiling experiments using the winsorization method with non-deduplicated data. We removed genes without polyA tails and kept genes with sufficient counts per million RPFs. After obtaining RPF counts from the coding regions for both ribosome profiling and RNA-seq, we performed CLR normalization and compositional linear regression, defining the residuals as TE for each gene in each sample. We averaged this sample-level TE based on cell lines and tissues. TEC is further calculated with rho scores[38]. To build an RNA co-expression matrix, we transformed CDS counts from RNA-seq experiments using CLR, averaged them based on cell lines and tissue, and calculated pairwise proportionalities (rho scores).

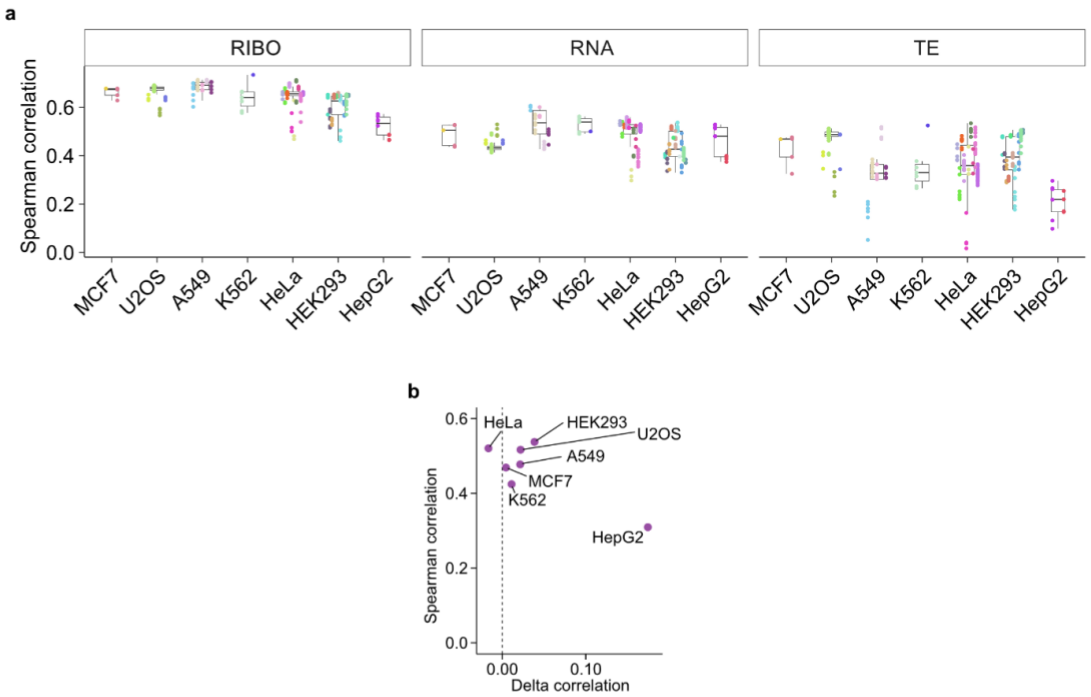


**Supplementary Fig. 8 | Spearman correlation between TE and protein abundance: a,** The correlation between protein abundance and clr-transformed RPF counts from ribosome profiling (left), clr-transformed read counts from RNA-seq (middle), or TE calculated with winsorized RPFs counts using the linear regression model (right). Individual dots indicate specific experiments colored according to study. In the boxplot, the horizontal line corresponds to the median. The box represents the IQR and the whiskers extend to 1.5 times of this range. **b,** TE was calculated with winsorized RPF counts without deduplication or with deduplication based on position and fragment length. The Spearman correlation coefficient between TE calculated with winsorized RPF counts and protein abundance[44] (y-axis) was plotted against "delta correlation" (x-axis) defined by subtracting the correlation values obtained with PCR deduplication from those obtained with the method using winsorized RPF counts without deduplication.

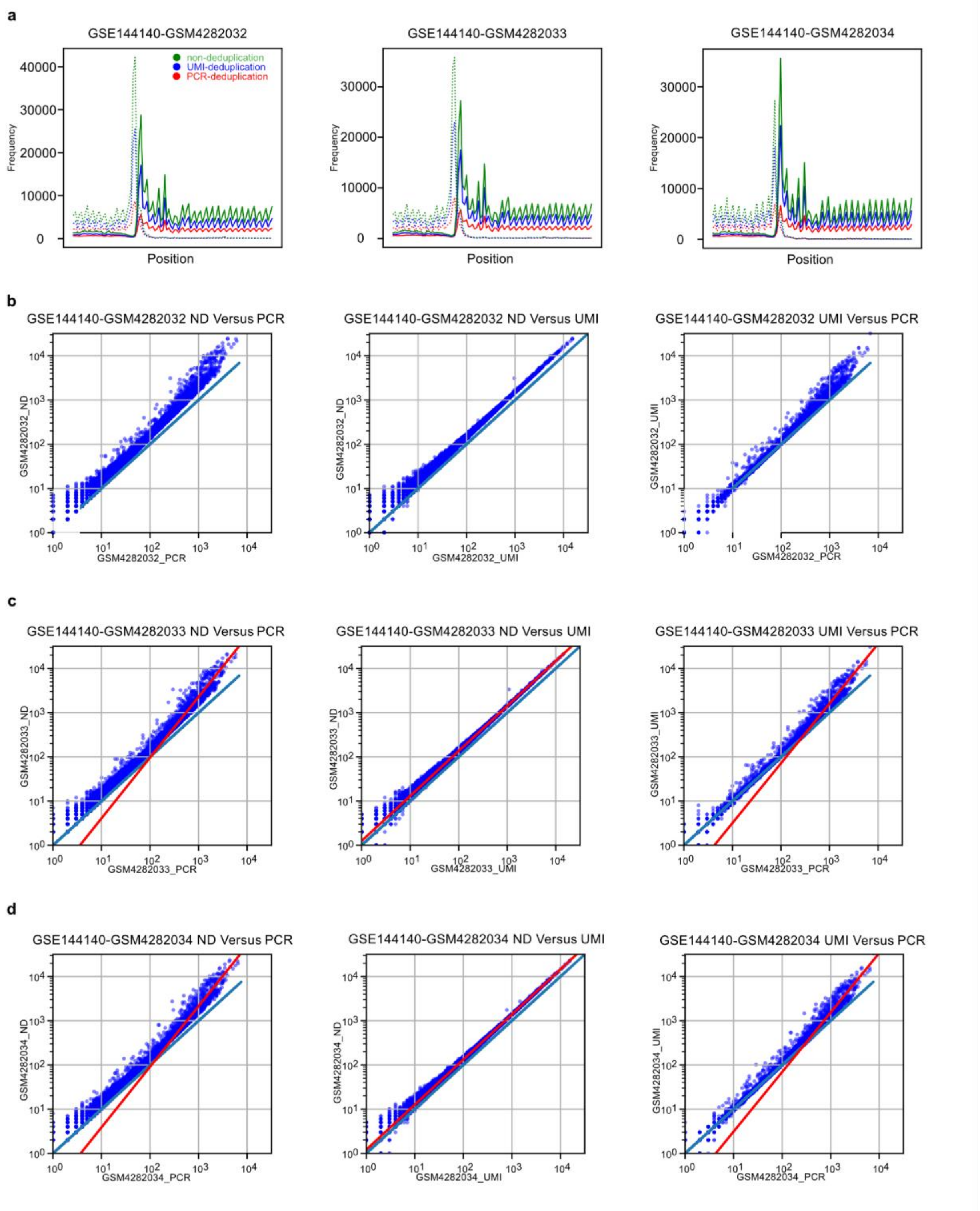


**Supplementary Fig. 9 | PCR vs. UMI deduplication comparison for GSE144140: a,** Metagene plots centered on the start codon for samples GSM4282032 (RPFs range: 28-36 nt), GSM4282033 (RPFs range: 28-36 nt range), and GSM4282034 (RPFs range: 26-35 nt range) were plotted using three different deduplication methods: non-deduplication (ND), UMI-deduplication (UMI), and PCR-deduplication (PCR). **b,** Correlation of gene counts for GSM4282032 between the three deduplication methods. A blue diagonal line represents a 1:1 ratio in all figure panels. Same analysis as panel b for GSM4282033 **c,** and GSM4282034 **d**.

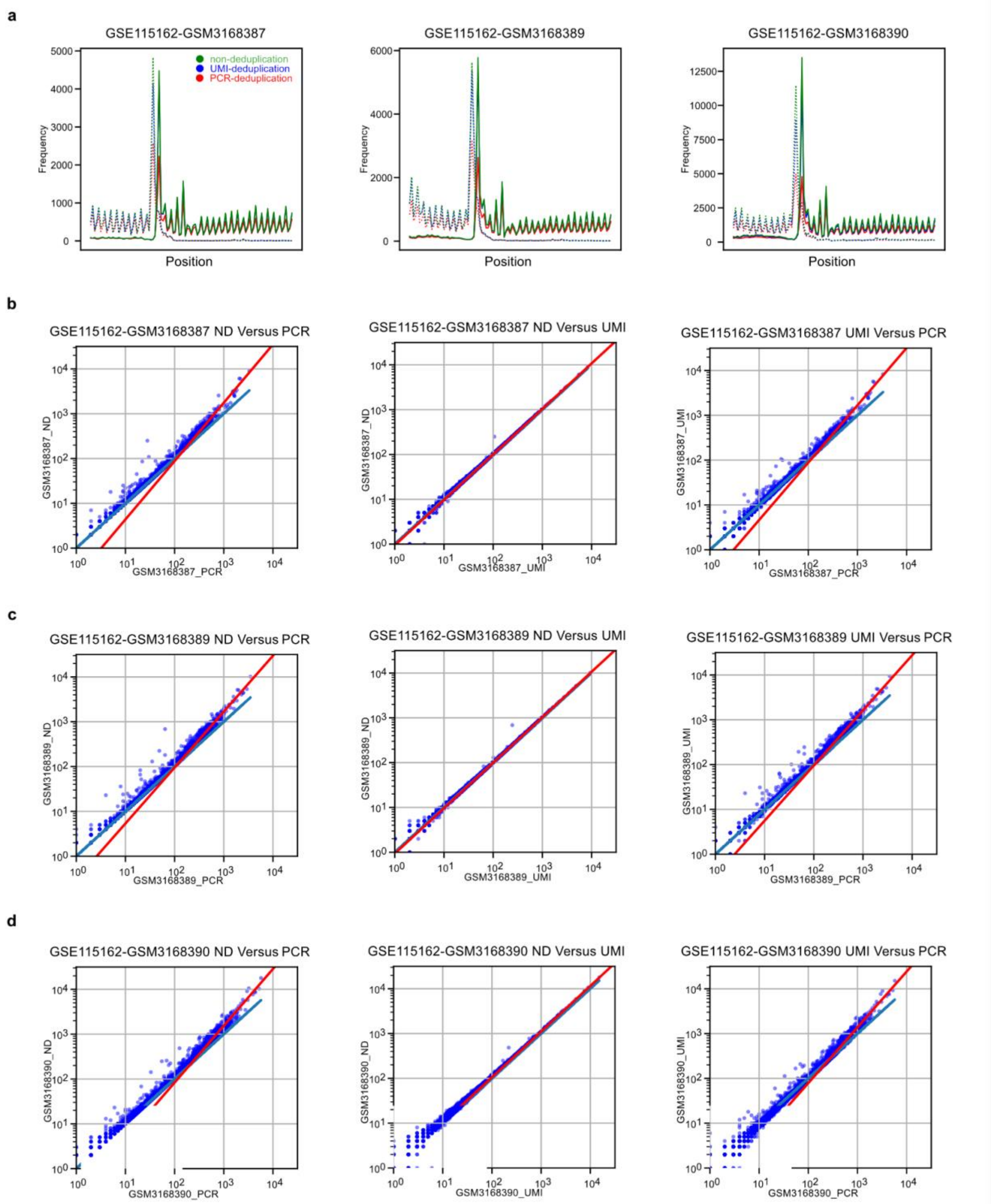


**Supplementary Fig. 10 | PCR vs. UMI deduplication comparison for GSE115162:** Similar analysis as Supplementary Fig. 7 for GSM3168387 (RPFs range: 24-34 nt), GSM3168389 (RPFs range: 23-33 nt), and GSM3168390 (RPFs range: 23-35 nt).

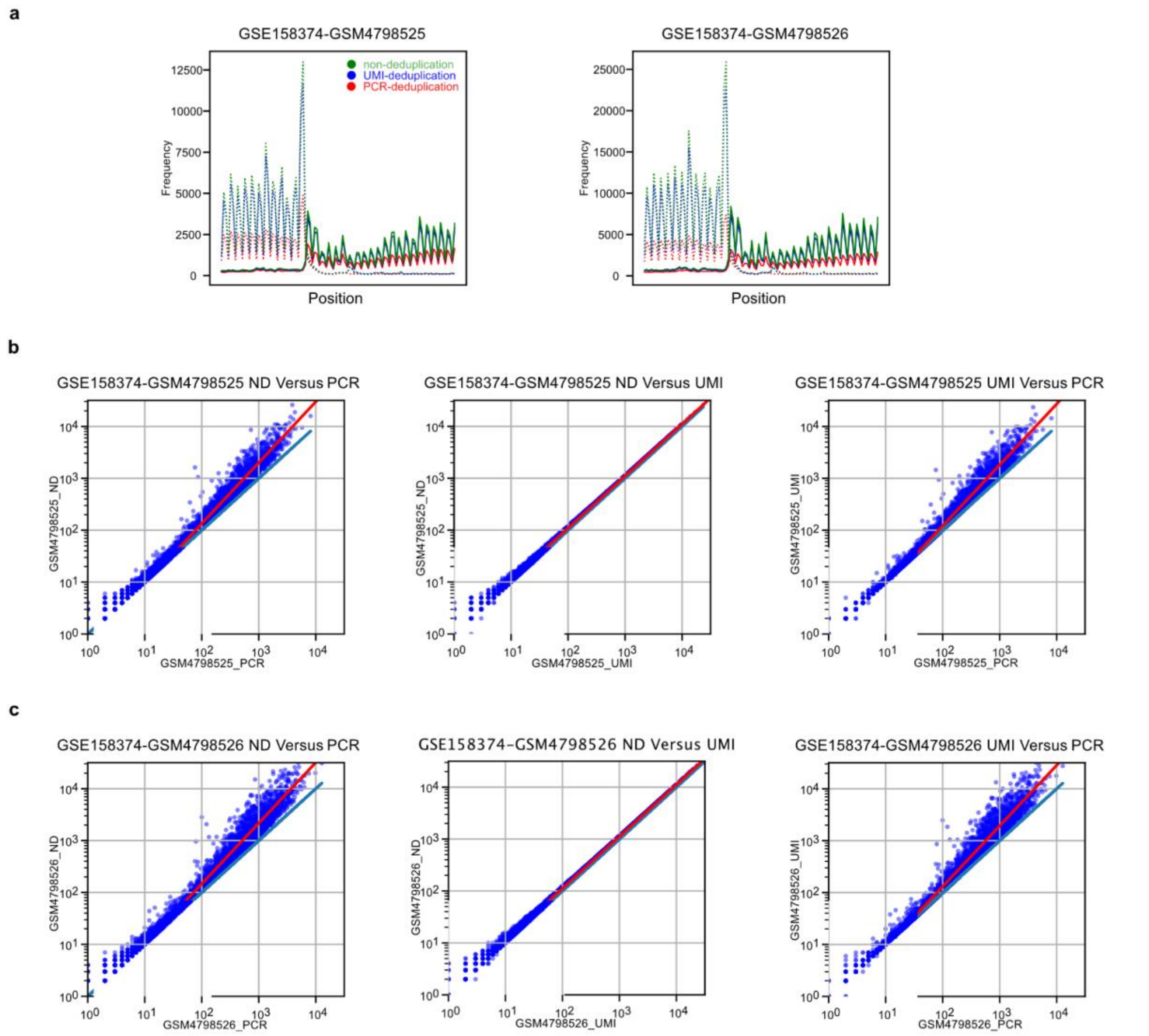


**Supplementary Fig. 11 | PCR vs. UMI deduplication comparison for GSE158374:** Similar analysis as figure S7 and S8 for GSM4798525 and GSM4798526, both in the 28-32 nt RPFs range.

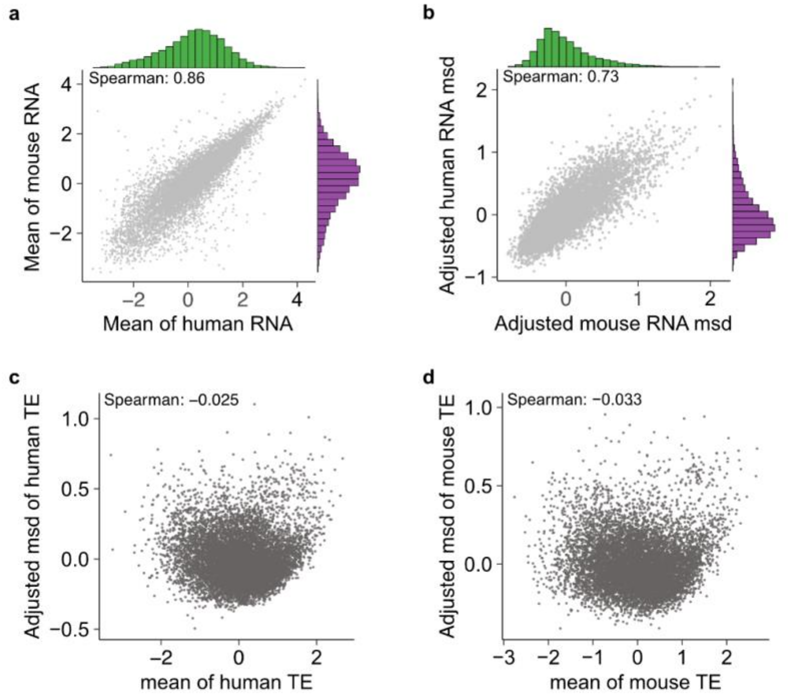


**Supplementary Fig. 12 | Conservation of gene expression between human and mouse: a,** The relationship between the mean RNA expressions (clr-transformed counts) of 9,194 orthologous genes across two species is plotted. Dots represent genes in all figure panels. **b,** The variability of genes' RNA expression was quantified with metric standard deviation (msd; Methods) across different cell lines and tissues in either human or mouse. To account for the correlation between mean RNA expression and its variability, we adjusted the msd values with their mean values (Methods). **c,** The scatter plot shows the adjusted msd values (y-axis; Methods) and the average TE across different cell types (x-axis) for human genes. **d,** Similar analysis as in panel c for mouse genes.

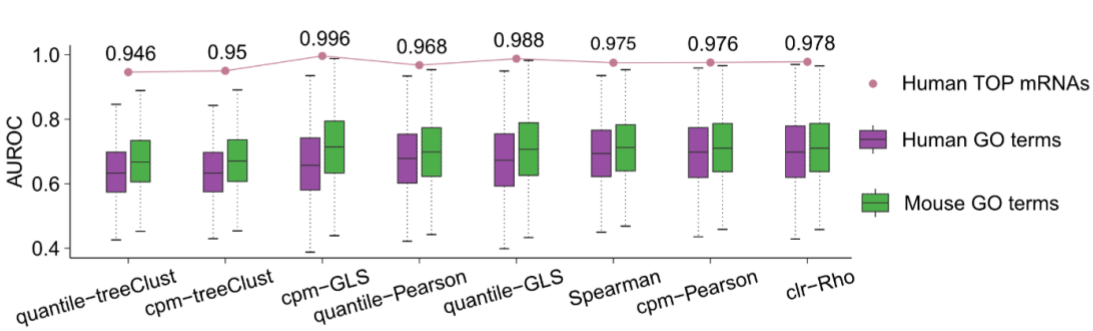


**Supplementary Fig. 13 | Evaluating the performance of eight methods to associate ribosome occupancy covariation with biological function: a,** The AUROCs for biological functions were calculated using the similarity scores among genes at ribosome occupancy level determined by eight distinct methods (Methods). In the boxplot, the horizontal line corresponds to the median. The box represents the IQR and the whiskers extend to the largest value within 1.5 times the IQR from the hinge. The dot in this figure represents the AUROC for human 5' TOP mRNAs.

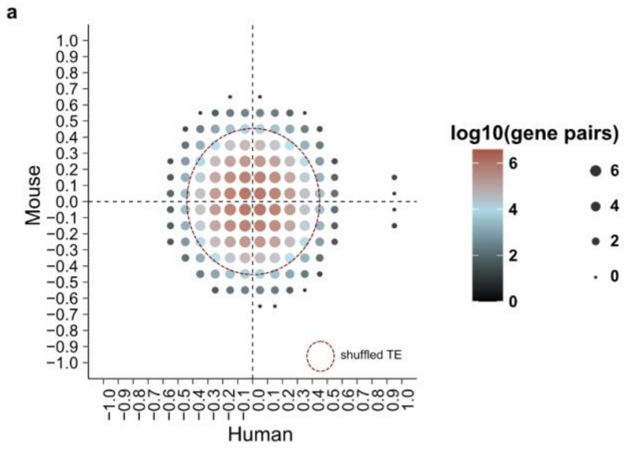


**Supplementary Fig. 14 | Lack of correlation in TEC across orthologous gene pairs between human and mouse using shuffled TE: a,** TE values that were randomly reassigned from the original data for each gene (shuffled) and TEC was calculated. In the figure panel, we plotted the number of orthologous gene pairs within specified ranges. Each dot represents the aggregated $\log_{10}$-transformed counts of these gene pairs. The dashed line captures 95% of the data.

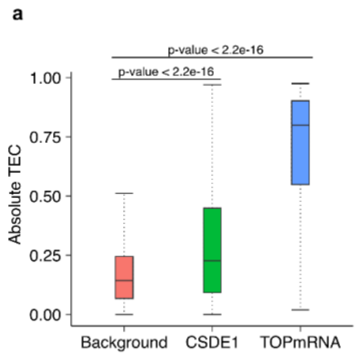


**Supplementary Fig. 15 | Comparison of TEC Distributions in TOP mRNAs and transcripts targeted by CSDE1: a**, Distribution of absolute TEC among 110 TOP motif-containing mRNAs[124] and 83 transcripts targeted by CSDE1 (table S23[48]) in comparison to all 11,149 human genes as background. Statistical significance between the groups was assessed using a Wilcoxon two-tailed test.

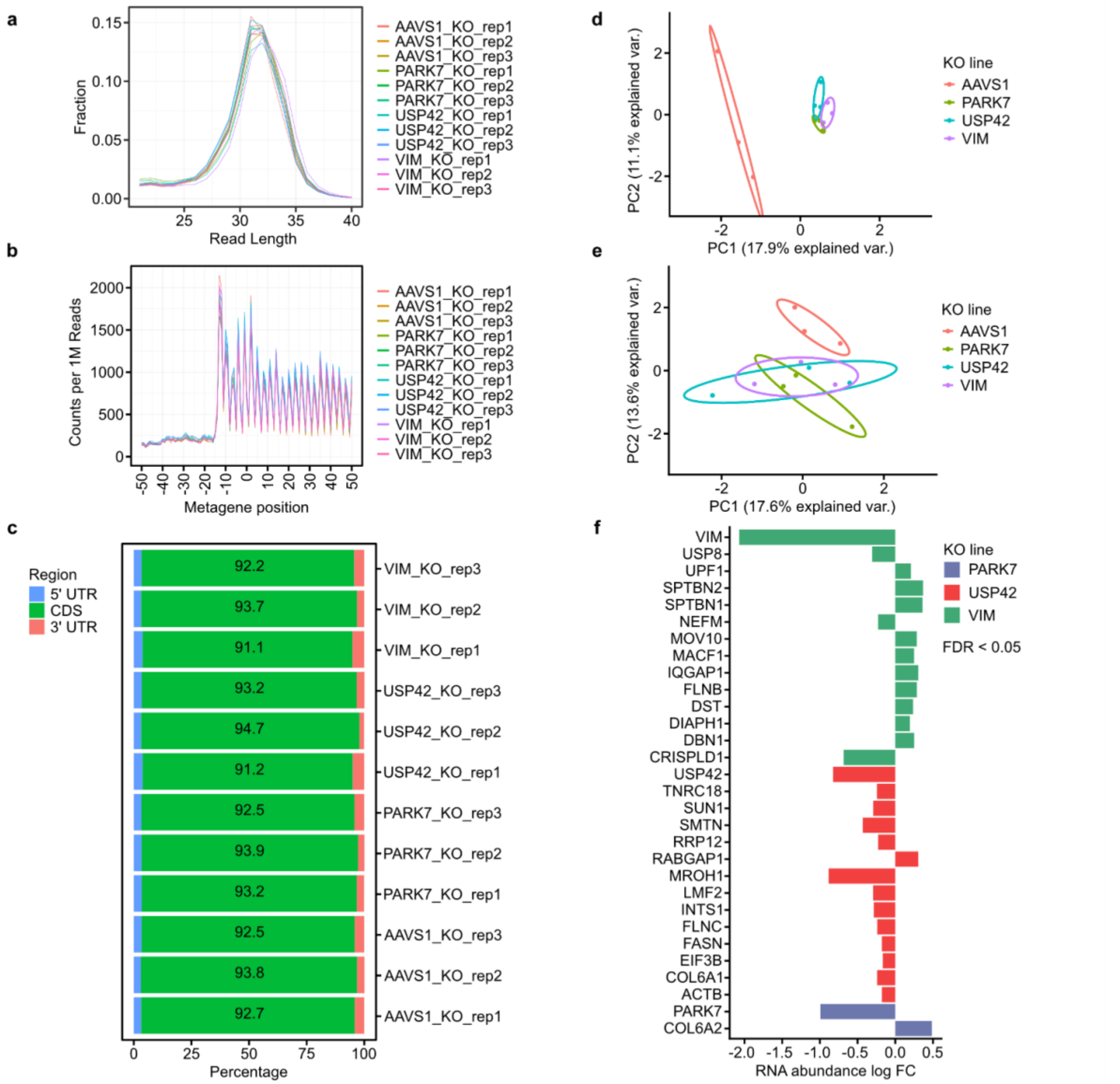


**Supplementary Fig. 16 | Ribosome profiling and RNA-seq of RBP KO cell lines:** For b through e, ribosome footprints between 28 and 35 nt were used. **a,** Read length distributions of ribosome footprints. **b,** Metagene plot at the start site. **c,** Location of mapped ribosome footprints. **d,** PCA was performed on standardized CPM reads for transcripts whose sum of CPMs across cell lines and replicates is in the top 80$^{th}$ percentile. PCA of RNA-seq counts. **e,** Same as D for ribosome profiling read counts. **f,** Differential RNA expression of KO cell lines. A significance threshold of FDR < 0.05 was used.

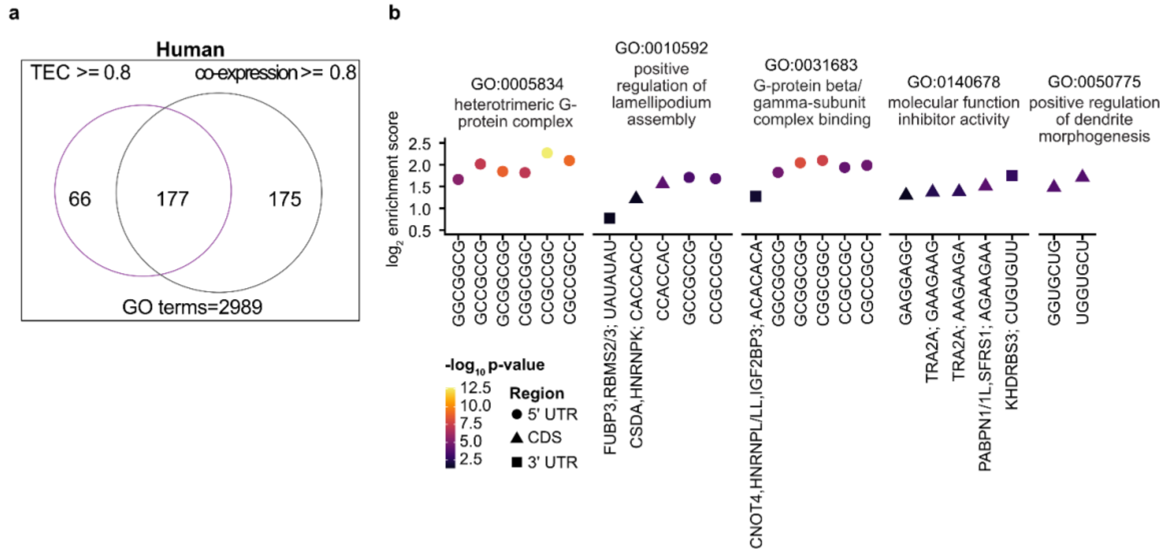


**Supplementary Fig. 17 | TEC and RNA co-expression among genes with shared functions in human: a,** A comparison between the number of human GO terms that have AUROC of 0.8 or higher with either TEC or RNA co-expression. **b,** Motif enrichment in human GO terms. RNA binding proteins (RBPs) from oRNAment[59] or Transite[60] are indicated. P-values were corrected using the Holm method and those kmers with a $p$-value $< 0.05$ are shown.

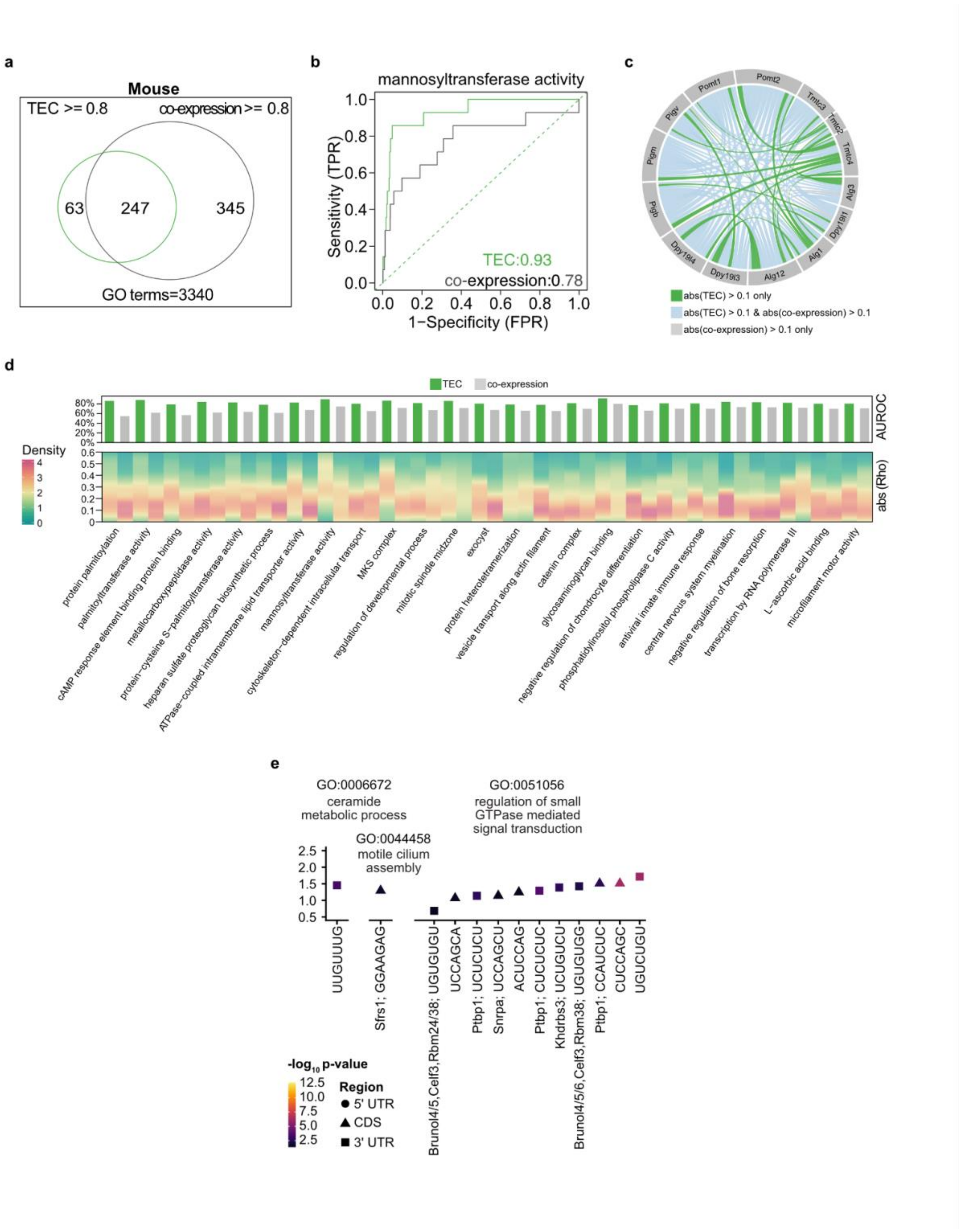


**Supplementary Fig. 18 | TEC and RNA co-expression among genes with shared functions in mouse: a,** Venn diagram for mouse GO terms that achieve an AUROC of 0.8 or higher with proportionality scores (rho) among genes at either TE or RNA expression level. **b,** The AUROC plot was calculated with genes associated with mannosyltransferase activity in mice. **c,** The connections represent absolute rho values above 0.1 in either TE pattern alone (green), in both RNA co-expression and TE pattern (blue), or RNA co-expression alone (gray). **d,** We summarized GO terms where genes exhibit greater similarity at the TE level than at the RNA expression level (AUROC with TEC > 0.8, and different AUROC between TEC and RNA co-expression > 0.1) in mice. We visualized the distribution of absolute rho score for gene pairs within each specific GO term (bottom; gene pairs with abs(rho) > 0.1) at the TE level. **e,** Motif enrichment in mouse GO terms. RNA binding proteins (RBPs) from oRNAment[59] or Transite[60] are indicated. P-values were corrected using the Holm method and those kmers with a p-value < 0.05 are shown.

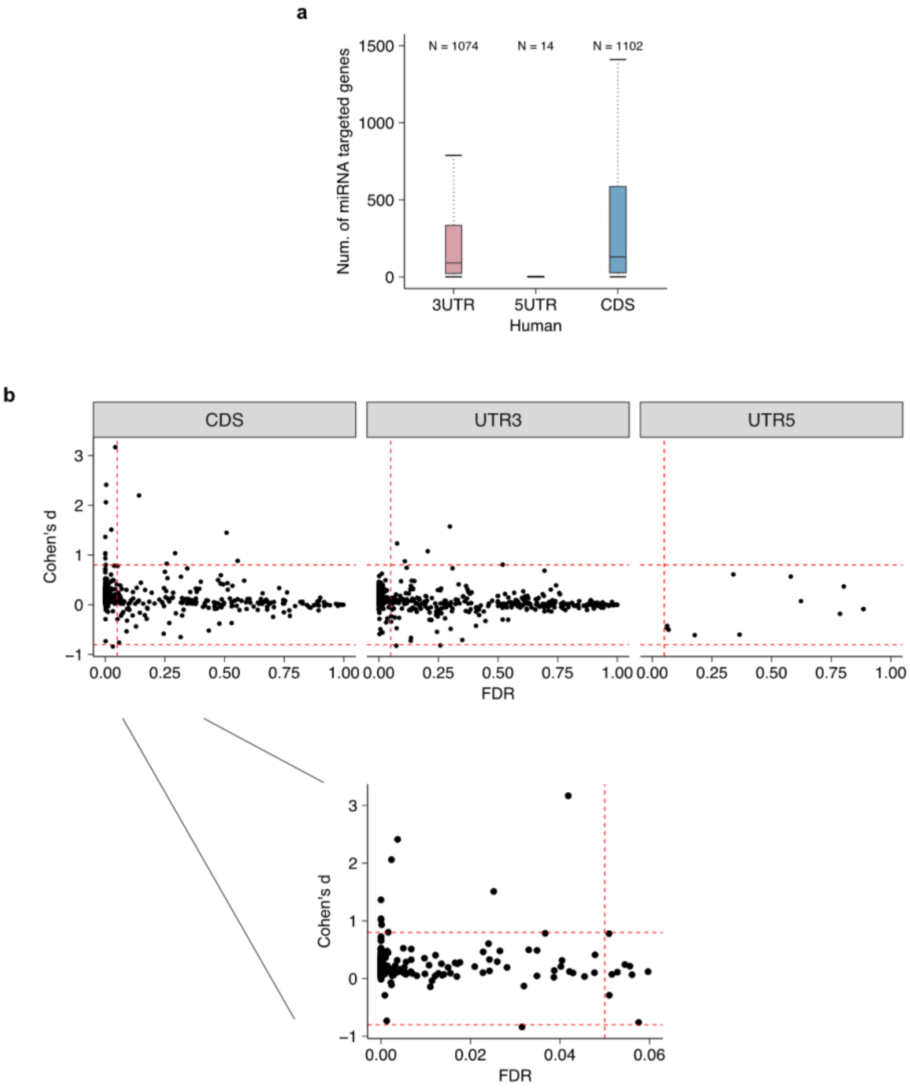


**Supplementary Fig. 19 | TEC is not primarily drive by shared miRNA binding sites:** Experimentally validated miRNA targets within 3' UTR, 5' UTR, and CDS were downloaded from https://dianalab.e-ce.uth.gr/tarbasev9[158]. 11,134 out of 11,149 human genes in RiboBase had at least one miRNA binding site. **a,** The distribution of the number of human genes targeted by miRNAs grouped according to transcript region. We only included miRNAs targeting at least 3 genes. ***N*** indicates the number of miRNAs in each boxplot. **b,** The distribution of TEC among 11,134 genes was used as the background. For each set of genes with a shared miRNA-binding site, the distribution of their TEC was compared to the background using a Student's t-test. Given the large number of genes in the background distribution, we incorporated both a FDR (<0.05) and an effect size threshold (Cohen's d > 0.8).

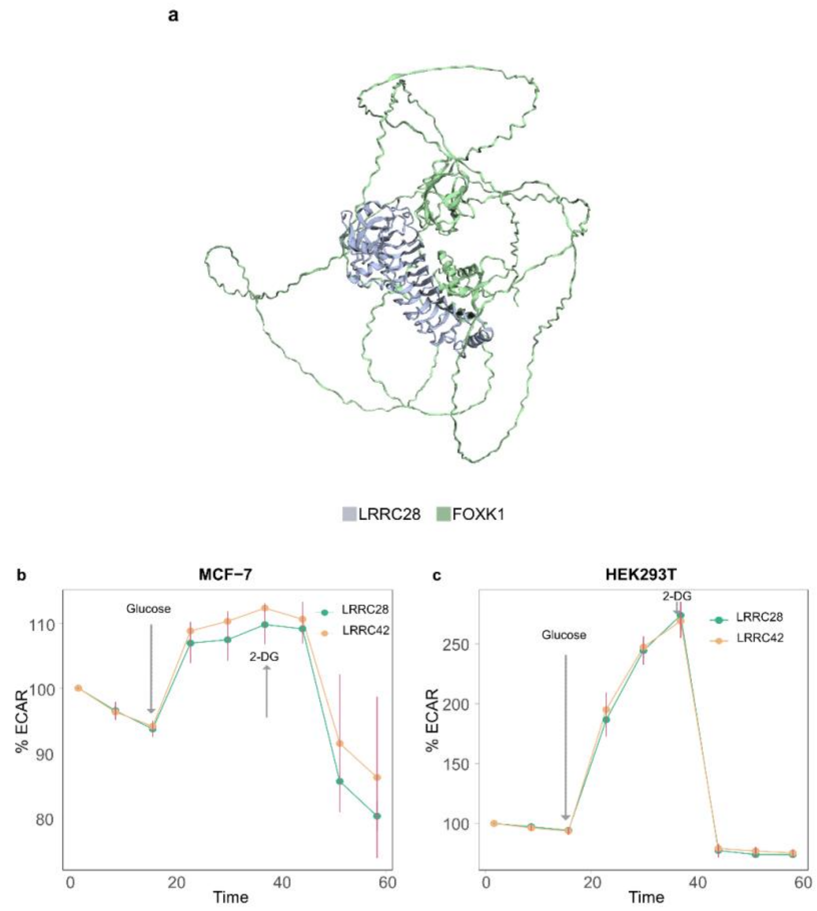


**Supplementary Fig. 20 | 3D structure of the interaction between LRRC28 with FOXK1: a,** AlphaFold2-multimer predicted binding between LRRC28 and FOXK1. Kinetic ECAR response of **b,** MCF-7 cell line (n=6, stable overexpression) and **c,** HEK293T cell line (n=6; stable overexpression) overexpressing LRRC28 or LRCC42 to 10 mM glucose and 100 mM 2-DG. Unpaired two-sided Student's *t*-test, ****P* < 0.005, ***P* < 0.05. Panel g & h shows mean ± s.d.; *n* shows biological independent experiments.

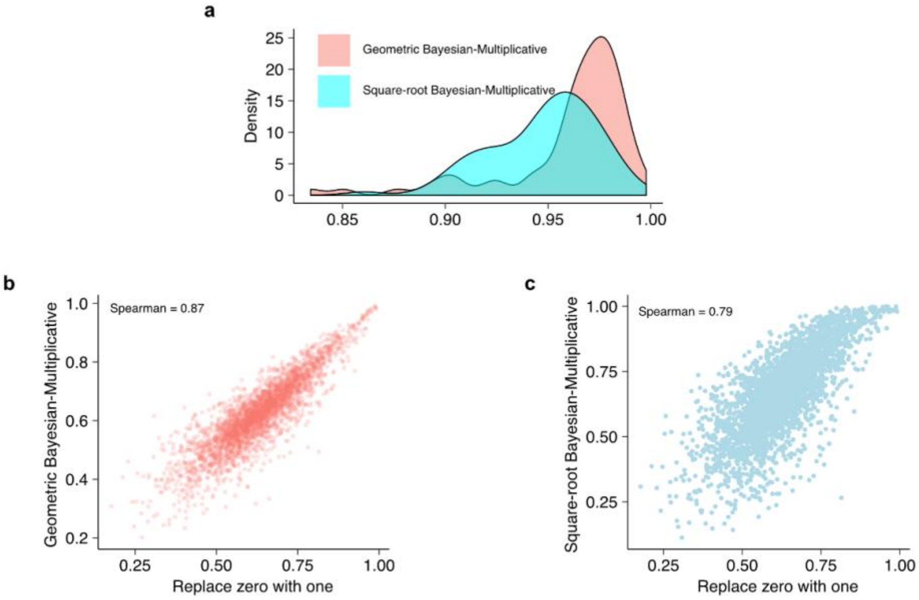


**Supplementary Fig. 21 | Zero imputation methods comparison: a,** Spearman correlation distribution for compositional TE across human cell lines and tissues comparing the current zero imputation method with two alternative methods. **b,** Spearman correlation of AUROC for biological functions between the current zero imputation method with Geometric Bayesian-Multiplicative method and **c,** Square-root Bayesian-Multiplicative method.

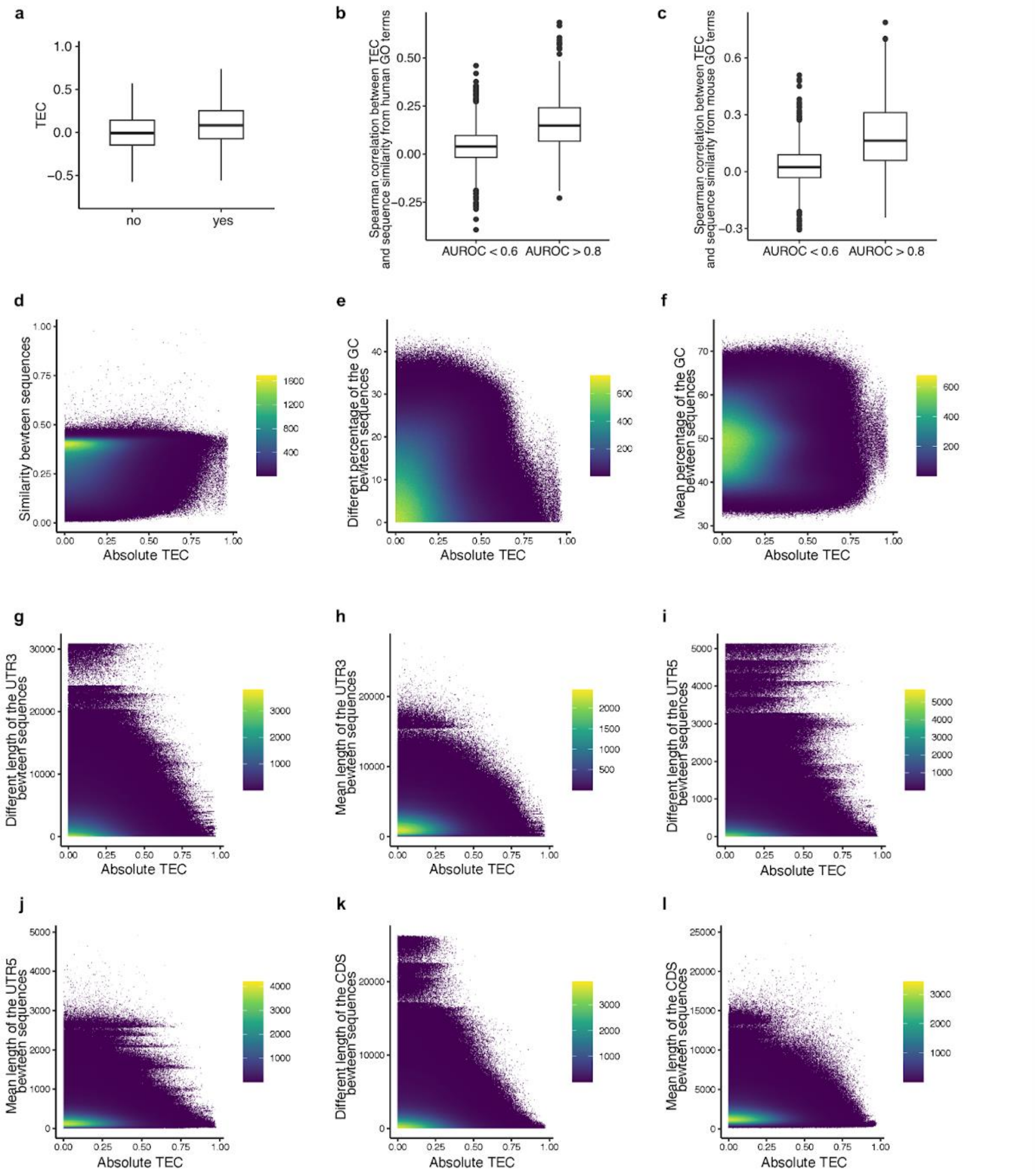


**Supplementary Fig. 22 | Correlations between genomic features and TEC: a,** TEC distribution of human gene pairs that either share the same protein domain or do not. **b,** Correlation between absolute TEC and sequence similarity for gene pairs within human biological functions; **c,** within mouse biological functions. **d,** Correlation between absolute TEC and sequence similarity among human gene pairs. **e,** Correlation between absolute TEC and the absolute difference in GC percentage between human gene pairs; **f,** and the mean GC percentage. **g,** Correlation between absolute TEC and the absolute difference in length of 3'UTR between human gene pairs; **h,** and the mean in length of 3'UTR. **i,** Correlation between absolute TEC and the absolute difference in length of 5'UTR between human gene pairs; **j,** and the mean in length of 5'UTR. **k,** Correlation between absolute TEC and the absolute difference in length of CDS between human gene pairs; **l,** and the mean in length of CDS. In figure panel d-l, we visualized the correlation between these features and TEC using a bin size of 1000.

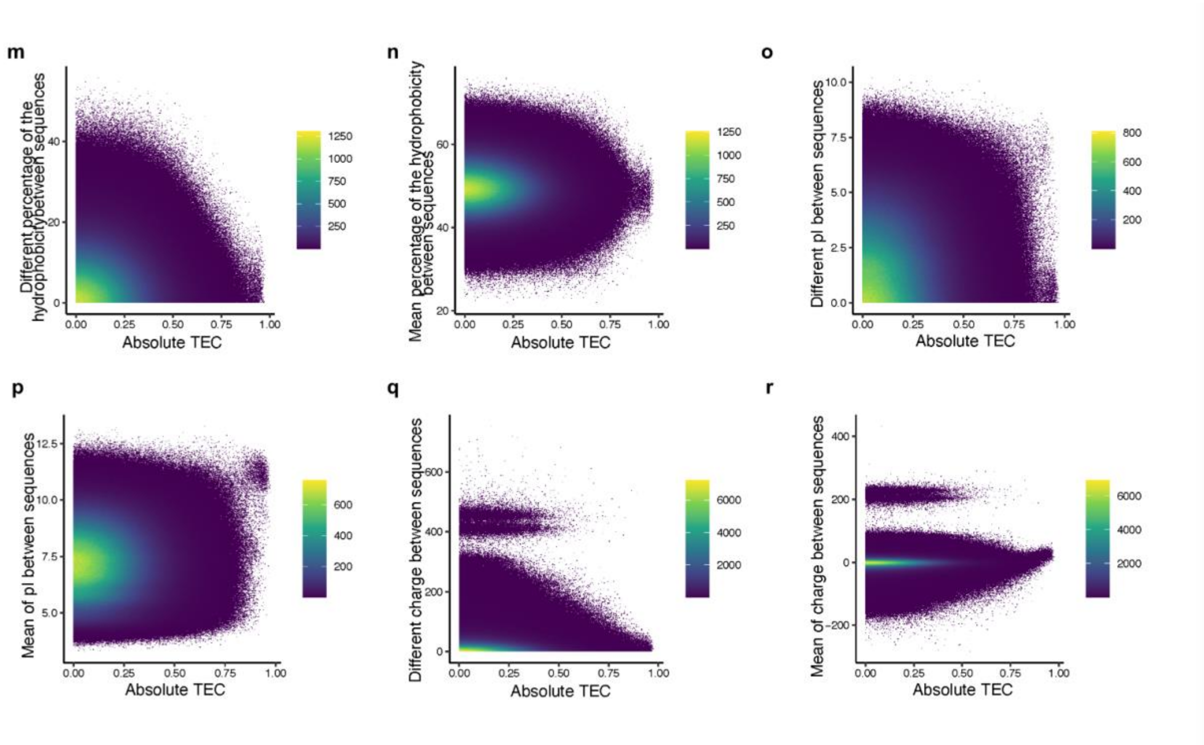


**Supplementary Fig. 23 | Correlations between protein features and TEC: a,** Correlation between absolute TEC and the different percentage of hydrophobic amino acids between human gene pairs; **b,** and the mean percentage of hydrophobic amino acids. **c,** Correlation between absolute TEC and the different pI between human gene pairs; **d,** and the mean of pI. **e,** Correlation between absolute TEC and the different charge between human gene pairs; **f,** and the mean charge. In this figure, we visualized the correlation between these features and TEC using a bin size of 1000.
